# The female (XX) and male (YY) genomes provide insights into the sex determination mechanism in spinach

**DOI:** 10.1101/2020.11.23.393710

**Authors:** Hongbing She, Zhiyuan Liu, Zhaosheng Xu, Helong Zhang, Feng Cheng, Xiaowu Wang, Wei Qian

**Author notes:** These authors contributed equally to this work (Hongbing She, Zhiyuan Liu). **Correspondence:** Wei Qian.

## Abstract

Sexual reproduction is the primary means of reproduction for the vast majority of macroscopic organisms, including almost all animals and plants. Sex chromosomes are predicted to play a central role in sexual dimorphism. Sex determination in spinach is controlled by a pair of sex chromosomes. However, the mechanisms of sex determination in spinach remain poorly understand. Here, we assembled the genomes of both a female (XX) and a male (YY) individual of spinach, and the genome sizes were 978 Mb with 28,320 predicted genes and 926 Mb with 26,537 predicted genes, respectively. Based on reported sex-linked markers, chromosomes 4 of the female and male genome were defined as the X and Y chromosomes, and a 10 Mb male-specific region of the Y chromosome (MSY) from approximately 95– 105 Mb, was identified that contains abundant transposable elements (92.32%). Importantly, a large-scale inversion of about 13 Mb in length was detected on the X chromosome, corresponding to ~9 Mb and ~4 Mb on the Y chromosome, which were located on both sides of the MSY with two distinct evolutionary strata. Almost all sex-linked/Y-specific markers were enriched on the inversions/MSY, suggesting that the flanked inversions might result in recombination suppression between the X and Y chromosomes to maintain the MSY. Forty-nine genes within the MSY had functional homologs elsewhere in the autosomal region, suggesting movement of genes onto the MSY. The X and Y chromosomes of spinach provide a valuable resource for investigating spinach sex chromosomes evolution from wild to cultivated spinach and also provide a broader understanding of the sex determination model in the Amaranthaceae family.

## Introduction

Hermaphroditism is a prevalent sexual system in flowering plants, while dioecy, constituting separate male and female individuals, is rare in angiosperms (Henry et al., 2018). Only ~6% of all angiosperms are dioecious, a trait which has independently evolved many times from hermaphroditic ancestors (Renner, 2014; Yang et al., 2020). A canonical theory called the two-mutation model was proposed to explain the evolution of dioecy. The model suggested that at least two tightly linked mutations were involved in the evolution of dioecy from hermaphroditism (Charlesworth and Charlesworth, 2018; Charlesworth, 2013). First, one recessive loss-of-function male-sterility mutation yielding female individuals occurred, which was followed by a dominant suppressor female function mutation resulting in female sterility and thus maleness. The suppression of recombination is an indispensable event for the evolution of dioecy, as it can maintain a stable dioecious population without cosexual or neuter individuals. The transposable elements or repetitive sequences accumulated increasingly within the two mutations as the non-recombining region. Finally, a Y-specific region is referred to as a male-specific region of Y chromosome (MSY) (Akagi et al., 2018; Charlesworth, 2019; Pannell and Gerchen, 2018).

Over the past decade, the advent of advanced next-generation sequencing (NGS) and third-generation sequencing (TGS) technologies, especially Pacific Biosciences (PacBio) and Oxford Nanopore Technology (ONT), has provided a possibility for uncovering the sex determination region and evolutionary history of sex chromosomes in dioecious plants. In papaya, the hermaphrodite-specific area of the Y^h^ chromosome (HSY) (8.1 Mb) is larger than its X counterpart (3.5 Mb), mostly due to retrotransposon insertions. Large-scale inversions are also likely to result in recombination suppression between the X and Y^h^ chromosome (Wang et al., 2012). In persimmon, a small RNA, *OGI*, was identified within the MSY (~1 Mb) that could determine the sex by targeting the autosomal *MeGI* gene (Akagi et al., 2014). Two genes, suppressor of female function (*SOFF*) and defective in tapetum development and function 1 (*TDF1*), were identified as regulating male and female function in asparagus (Harkess et al., 2017). Similar to the observation in asparagus, two Y-specific genes, Shy Girl (*SyGl*), suppressing female function, and Friendly Boy (*FyBy*), accelerating male function, were also identified in kiwifruit (Akagi et al., 2018; Akagi et al., 2019). In addition, the repeated translocation of a female-specific sequence (13 kb) is able to drive sex-chromosome turnover in strawberries (Tennessen et al., 2018). Although the two-mutation model is a plausible one, it may not be the only path to dioecy, as mentioned above. Taken together, sex determination mechanisms are determined by many factors, and thus understanding the evolutionary history of sex chromosomes or unraveling sex determination mechanisms in plants requires further investigation.

Spinach (*Spinacia oleracea* L.) is an important leafy vegetable crop belonging to the family Amaranthaceae (Cai et al., 2018). Spinach is mostly dioecious with separate male and female individuals, while occasional monoecy with both male and female flowers is observed (Deng et al., 2013). The gender of spinach is controlled by an allelic pair named X and Y, which are located at the largest chromosome (Iizuka and Janick, 1962). Spinach has the XY sex-determining system, with no obvious size difference between the X and Y chromosome (Deng et al., 2013). Previous studies developed several sex-linked markers, including SO4 (Groben and Wricke, 1998), D4.3(Liu et al., 2015), and 12 fully sex-linked single nucleotide polymorphisms (SNPs), which were validated in large-scale populations (>1400) of over 100 spinach germplasm accessions and cultivars (Okazaki et al., 2019). Additionally, male-specific markers were also identified: T11A, V20A (Akamatus et al., 1998), S5.7, and S9.5 (Liu et al., 2015). Based on the male-specific markers, five bacterial artificial chromosomes (BACs) were cloned with a total length of ~690 kb, encompassing large amounts of repetitive elements; however, no sex determination genes were found (Kudoh et al., 2017). Recently, She et al. (2020) identified a male-specific region (~21 Kb) on chromosome 4 where the females had a low reads coverage, while the males had high coverage. One SNP named SponR, which was closely linked to the region, was developed that could be used for discriminating the XX, XY, and YY genotype (accepted). By constructing two high-density genetic maps, Yu et al. (2020) identified a non-recombining region with 39 bin markers co-segregating with sex, which were located at 45.2 cM of LG1. The region contained higher percentages of repetitive sequences and low gene density, and its X counterpart was estimated to be approximately 18.4 Mb (Yu et al., 2020). A high-quality reference genome is a powerful tool for gene mapping, evolutionary analysis, and comparative genomics. Xu et al. (2017) published the first chromosome-level spinach genome, but only 47% of sequences were anchored on chromosomes (Xu et al., 2017). Owing to the lack of a high-quality reference genome, the sex determination region remains unclear in spinach. Specifically, the vast majority of sex-linked markers are located at different scaffolds due to the incomplete genome; for instance, T11A located at Super_scaffold_113, D4.3 located at SpoScf_02571, and even the non-recombining region identified by Yu et al. encompassed various scaffolds or fragments of chromosomes (Yu et al., 2020).

To define the complete sex determination region, understand its origin, and authentically estimate the divergence time between the sex determination region and its X counterpart, an assembly of the X and Y chromosomes is urgently needed. A major obstacle to the assembly of the Y chromosome relates to the separation of the Y chromosome from male (XY) individuals. In this study, we found a male (YY) individual and report a high-quality genome assembly of female (XX) and male (YY) spinach using ONT technology and a high-density genetic linkage map. A comparison of the X and Y chromosome revealed the sex determination region in spinach, which was confirmed by two independent approaches. Furthermore, a large-scale inversion might suppress recombination between the X and Y chromosome, generating a stabilized MSY.

## Results

### Female (XX) and male (YY) genome assembly and annotation

A female (XX) and male (YY) individual from the inbred line 10S15 were used for genome assembly using ONT platform. Analysis of the 21-mer sequence of the Illumina reads (Table S1) revealed that the heterozygosity of the female and male was 0.076% and 0.119%, respectively. For the female individual, a total of 58.28 Gb of Oxford Nanopore reads, achieving ~58-fold coverage of the spinach genome, were assembled using Nextdenovo (v2.2.0), resulting in a 975 Mb assembly with a contig N50 size of 34 Mb (Table 1). Using a similar assembly strategy, the male individual genome was assembled using 42.06 Gb Oxford Nanopore reads, resulting in a 926 Mb assembly with a contig N50 size of 41 Mb (Table 1).

**Table 1.**
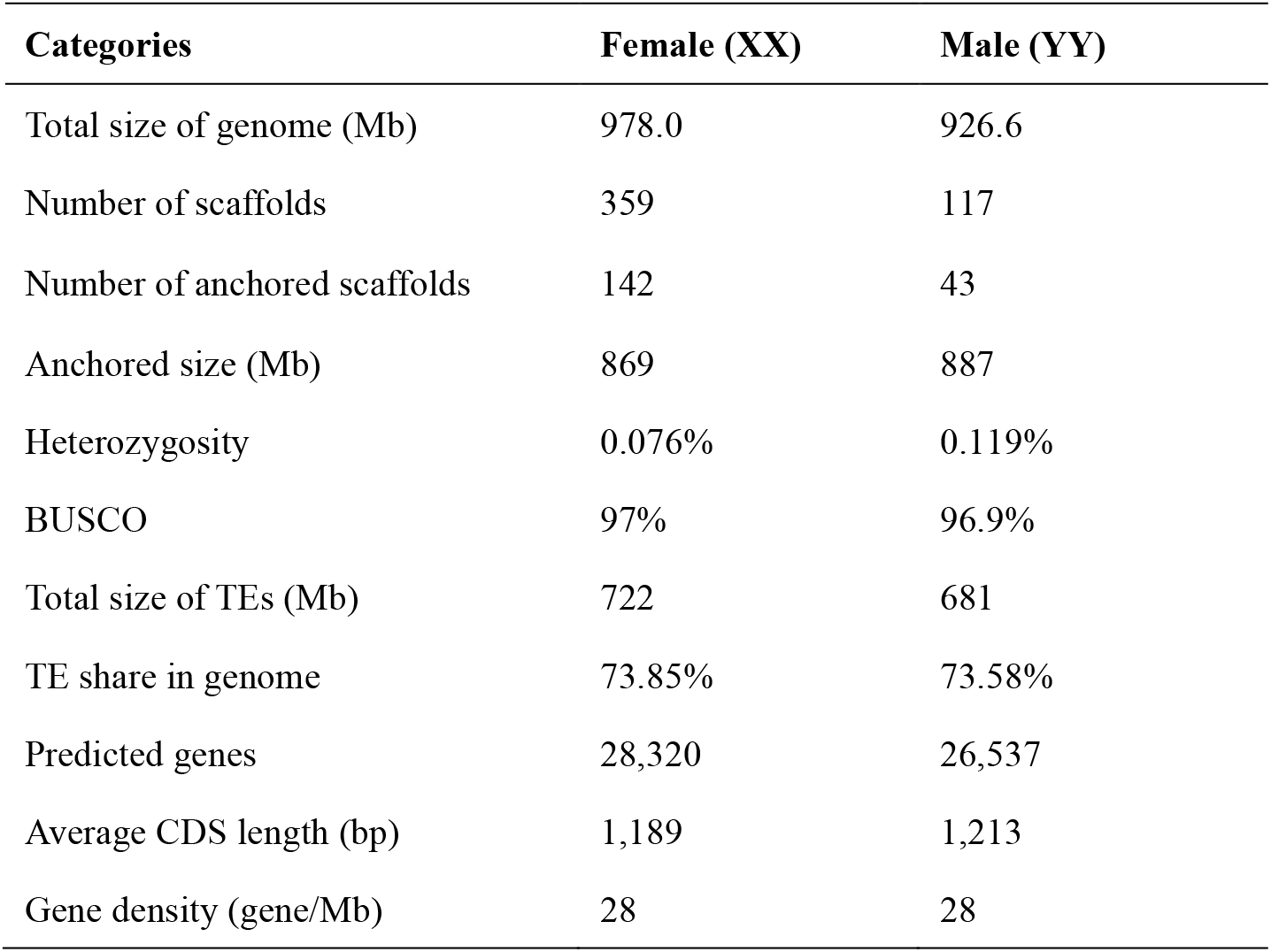
Characteristics of the spinach genome assembly.

Previously, we constructed a high-density genetic linkage map using a BC_1_ population that was used to anchor the contigs to pseudochromosomes (Qian et al., 2017). Sequence alignment by Blast with the initial assembled contigs from both the female and male genomes, and ordering of the specific-length amplified fragment (SLAF) markers in the linkage map, facilitated the ordering and orientation of the contigs. Additionally, the female and male genomes were also assembled by NECAT (v0.01) (Table S2) to correct and extend the contigs when conflict was occurred. For example, a contig199 from the XX genome assembled by NECAT conflicted with the XX genome assembled by Nextdenovo (Figure S1A). Sequence alignment between contig199 and the linkage map indicated that the contig was mis-joined, and it was corrected and extended based on the homolog region (Figure S1B). At last, the extended contig was obviously mapped on linkage group 4 (LG4) (Figure S1C). The integrated approach anchored 142 contigs to six pseudochromosomes for the female, comprising 88% (869 Mb) of the female genome assembly (Sp_XX_v1). The male genome also contained six pseudochromosomes of 887 Mb in length, encompassing 95.7% of the total assembly (Sp_YY_v1) (Table 1 and Figure S2).

The quality of the two genomes was assessed by four independent methods. First, the Illumina reads of six individuals were mapped to the female and male assembly using Burrows-Wheeler Aligner (BWA) (v0.7.17) (Li and Durbin, 2009), which shared perfect alignments with proper mapping rates of >92.31% (Table S3). Second, approximately 97% of embryophyte genes were detected in both the female and male assembly by Benchmarking Universal Single-Copy Orthologs (BUSCO) (Table 1 and Table S4) (Waterhouse et al., 2017). Next, sequence alignment of both genomes indicated they had high sequence identity and good alignment, except for several inversions located on Chr4 (Figure S3). Finally, 126.81 Gb of Hi-C clean data with 126× coverage generated from a male individual of an inbred line 12S4 (unpublished) were aligned to the female and male assembly to assess their quality (Figure S4).

The both genomes had 73.85% and 73.58% repetitive sequences for female and male, respectively. The majority of repetitive sequences in our assembly were long terminal repeat (LTR) retrotransposons, especially Gypsy-LTRs and Copia-LTRs (Table S5). Using a combination of *ab initio*, protein-homology-based, and RNA-seq-based methods (Table S6), we predicted 28,320 and 26,537 protein-encoding genes in the female and male genomes, respectively, and approximately 90% genes were successfully annotated (Table S7 and Table 1). In addition to protein-coding genes, we identified 2817 tRNAs, 289 rRNAs, and 718 miRNAs in the male genome, while 3037 tRNAs, 471 rRNAs, and 800 miRNAs were identified in the female genome. The Circus tool was used to illustrate the repeat density, gene density and syntenic relationship between the female and male genome (Figure 1). Comparative genomics confirmed that the spinach was relatively closely related to *C. quinoa* and then *B. vulgaris* (Figure S5A and Table S8). The syntenic analysis of the three species strongly indicated that the genome of the species shared many syntenic regions, suggesting a high level of chromosome rearrangements among the three species during their evolution (Figure S5B).

**Figure 1.**
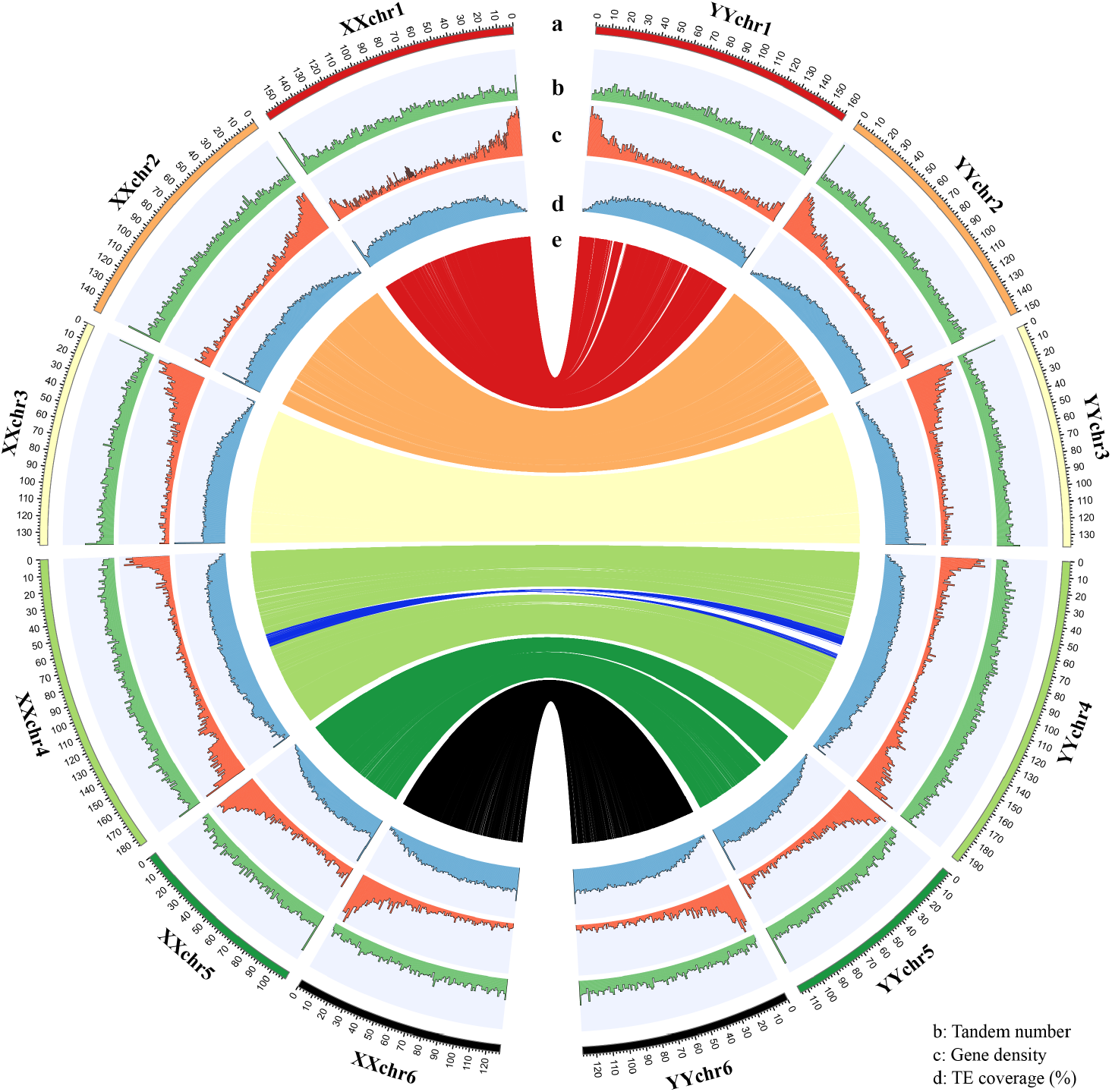
Genomic landscape between the female and male genome. **(a)** Ideogram of the chromosome from the female and male genome (in Mb scale). **(b)** Tandem number represented as the number of tandem repeats per 20 Kb. **(c)** Gene density in 20-Kb windows. **(d)** Percentage of coverage of transposable elements per 20-Kb. **(e)** Syntenic gene pairs between the female and male assemblies. Blue lines represent sex determination region

### Identification of a male-specific genomic region

To identify the sex determination region, we sought to map the reported sex-linked/Y-specific markers against the female (XX) and male (YY) genomes, respectively. Interestingly, almost all of the markers were enriched on XXchr4 (from 85.2 to 98.3 Mb) and YYchr4 (from 86.1 to 109 Mb), and thus XXchr4 and YYchr4 were defined as the X chromosome and Y chromosome in spinach, respectively, and the region on the sex chromosomes was considered as a potential sex-determining region (Figure 2).

**Figure 2.**
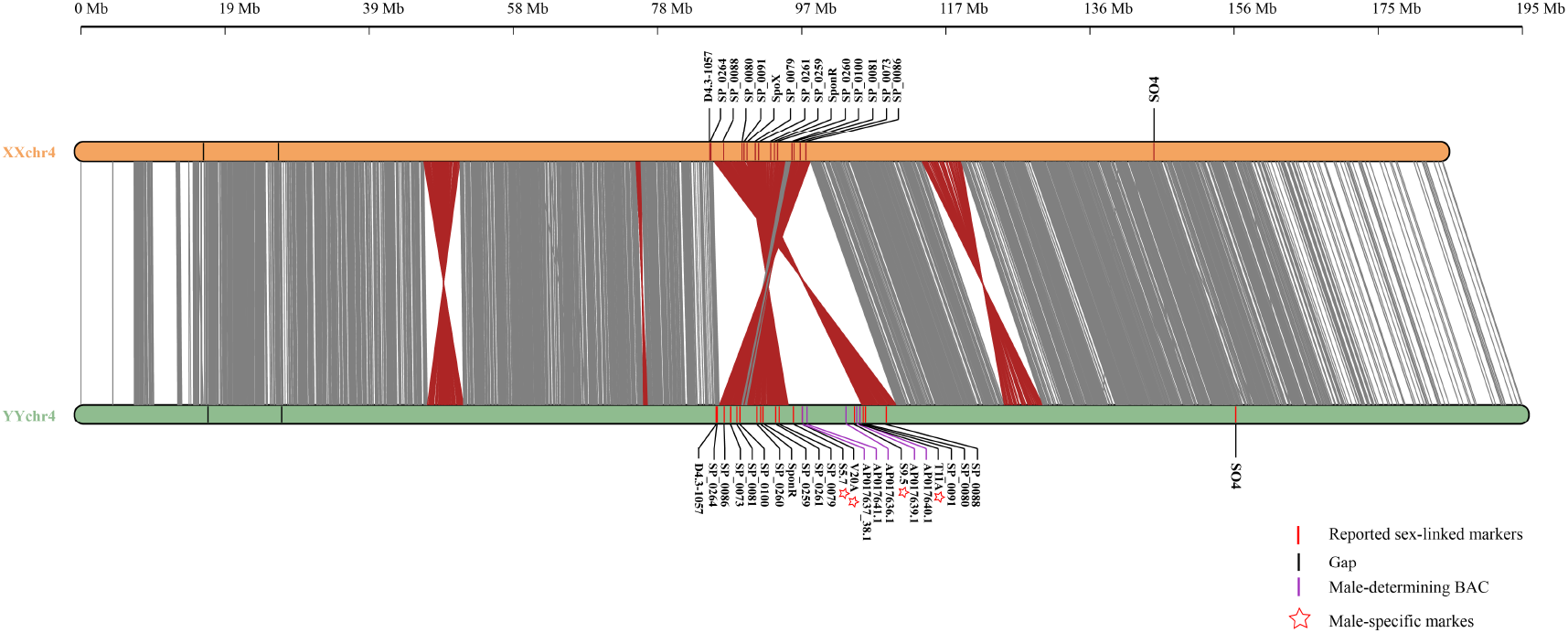
Sequence alignment of the XXchr4 (X) and YYchr4 (Y) chromosome. The grey and red lines between the X and Y chromosome indicate noninverted and inverted alignments, respectively. Reported sex-related markers were mapped to sex chromosomes, and the name is shown on the top/bottom of the chromosome.

Meanwhile, sequence alignment of the X and Y chromosome indicated the presence of three large-scale inversions, which was confirmed using the female and male ONT long reads (Figure S6, Figure S7 and Figure S8). Most importantly, the second inversion (IV2) was located on the sex-determining region, and thus IV2 might be related to sex determination in spinach. Based on the position of IV2 on the Y chromosome, the IV2 was further divided into two parts, IV2-1 from 90.3–98.8 Mb on the X chromosome, corresponding to 86.5–95.8 Mb on the Y chromosome, and IV2-2 from 85.5–90.1 Mb on the X chromosome, corresponding to 105.9–110.6 Mb on the Y chromosome. Interestingly, IV2-1 and IV2-2 were located on both sides of a 10 Mb region on the Y chromosome, which was defined as a potential MSY, as it did not have a counterpart on the X chromosome and contained male-specific markers (T11A, V20A, S5.7, and S9.5) and male determination BAC sequences (Figure 2). Recombination suppression is a vital event during sex chromosome evolution, as it facilitates a stable dioecious population. Therefore, the large-scale IV2-1 and IV2-2 flanked by the MSY might suppress recombination between the X and Y chromosome and maintain the conservation of MSY, thus generating separate female and male individuals in spinach.

Additionally, the MSY was further determined by four independent approaches as follows: (1) assessment of alignment by mapping the female genome (Dohm et al., 2013) to the Y chromosome (Figure S9A); (2) examining the mapped reads count when aligning 20 male versus 20 female resequencing reads against the Y chromosome (Table S1 and Figure S9B); (3) identification of fully sex-linked SNPs and InDels by aligning 59 individual resequencing reads to the YY genome (Table S1 and Figure S9C); and (4) examining the recombination rate by mapping SLAF markers to the Y chromosome (Figure S9D). Taken together, the MSY identified above was confirmed as a male-specific region with suppression of recombination and female coverage. In addition to the reported sex-linked markers, 2,227 fully sex-linked SNPs and 66 fully sex-linked InDels within the 59 individuals were enriched on IV2 (Figure 2 and Figure S9C), which is flanked by the MSY. The structure of the MSY is consistent with other species, such as persimmon and papaya (Akagi et al., 2014; Wang et al., 2012).

### Validation of the sex-determining region using two independent approaches

To further verify the accuracy of the sex-determining region above, we applied two independent approaches, including a genome-wide association study (GWAS) using the spinach v1 genome as a reference (Xu et al., 2017) and a reference-blind approach (Akagi et al., 2018).

First, 32 female and 48 male individuals (Table S1) were used to GWAS and characterize the sex determination region using spinach v1 (Xu et al., 2017) as the reference genome. A total of 1,665 SNPs were detected as significantly associated with sex (α<0.05; Table S9). Among these SNPs, approximately 40.6% of SNPs were located at the distal end of Chr3 and Chr4 from 54.5–59.2 Mb, and 59% SNPs were mapped at scaffolds due to the incompleteness or inaccuracy of the spinach v1 genome assembly. For the associated SNPs in scaffolds, more than 59.5% of SNPs were located on Super_scaffold_232, Super_scaffold_60, and Super_scaffold_66 (Figure S10A, C). Additionally, the vast majority of genotypes (>99%) of these sex-associated SNPs were homozygous in females, while more than 94% of the genotypes were heterozygous in males (Table S9 and Figure S10C), demonstrating that an XY system is involved in sex determination in spinach.

Next, we compared the alignment free k-mer counts of 15 male and 16 female sequence pools from inbred lines to identify the sex-determining region (Table S1). A total of 461,190 male-specific k-mers (31-bp) (MSKs), which were completely absent from the female reads were identified. Paired-end reads, including MSKs, were used to assemble 4,659 potential initial contigs, and these male-specific reads were mainly enriched on Chr3 (112–113 Mb) and Chr4 (55.4–59 Mb) of the spinach v1 reference genome (Xu et al., 2017) (Figure S10D). Based on the how the SNPs within each of these contigs segregated with sex in the 31 individuals, 1,362 sex-linked contigs and 304 Y-specific contigs were identified from 4,659 initial contigs. As for the 1,362 sex-linked contigs, 64.7% contigs were mapped at scaffolds (the top three scaffolds were Super_scaffold_232, Super_scaffold_60, and Super_scaffold_66), and approximately 21.8% of sex-linked contigs were located on Chr4 and were enriched from 55.4 to 59 Mb, followed by Chr3, the region of which were mainly mapped at the distal end of Chr3 of the spinach v1 genome (Figure S10E). All of the above results are consistent with the GWAS, consistently indicating that the five major sex-linked regions identified in the study are reliable regions that could be involved in sex determination in spinach. Unfortunately, the 304 Y-specific contigs were not significantly enriched at a specific region of the spinach v1 genome. This might be due to the incompleteness or inaccuracy of the spinach v1 genome assembly.

Therefore, we further examined the sex-related sequences identified by GWAS and k-mer using the male (YY) genome. The male-specific reads identified above were mainly mapped on the YYchr4 chromosome from 86–110.6 Mb (Figure S11A). 93.4% of Y-specific contigs and 70.9% of sex-linked contigs could be mapped on the Y chromosome from 86.1 to 110.5 Mb, which is the region of IV2 and MSY (Figure S11B and Table S10 and Table S11). Importantly, the five main sex-associated regions identified above were mapped on the Y chromosome from 86.5–95.8 Mb and 105.9–110.6 MB, corresponding to IV2-1 and IV2-2, respectively (Figure S11C), greatly demonstrating the authenticity of sex-determining region identified above. Additionally, this is further indicative of the higher continuity and completeness of the genome assembled in this study than in the previous spinach v1 genome, as the distal end of the Chr3 might be a mis-join (Figure S11D) (Xu et al., 2017).

### Analysis of repetitive sequences and nucleotide substitution rates

Previous investigations proposed that the sex determination region is full of repetitive sequences and possesses low gene density (Jia et al., 2019; Wang et al., 2012). In this study, the sex determination region on the Y chromosome exhibited greater amounts of transposable elements and tandem repeats but lower gene density relative to its counterpart on the X chromosome (Figure 3). Specifically, the sex-determining region of the Y chromosome from 86.5–110.6 Mb and its X counterpart from 85.8–98.8 Mb shared 90.44% and 88.47% repetitive sequences, respectively, which is much higher than the spinach-genome-wide average of ~73% (Table 1 and Table S12). LTR/Copia elements are the most abundant repeats in both the sex determination region on the Y chromosome and its X counterpart. The IV2 was further divided into two parts, IV2-1 (86.5–98.8 Mb) and IV2-2 (105.9–110.6 Mb), according to the physical position of MSY on the Y chromosome (Figure 3). Comparison of transposable elements on IV2-1 and IV2-2 revealed that IV2-2 accumulated more transposable elements than IV2-1 on both the X and Y chromosome, and MSY contained the highest transposable elements (92.32%) (Table S13).

**Figure 3.**
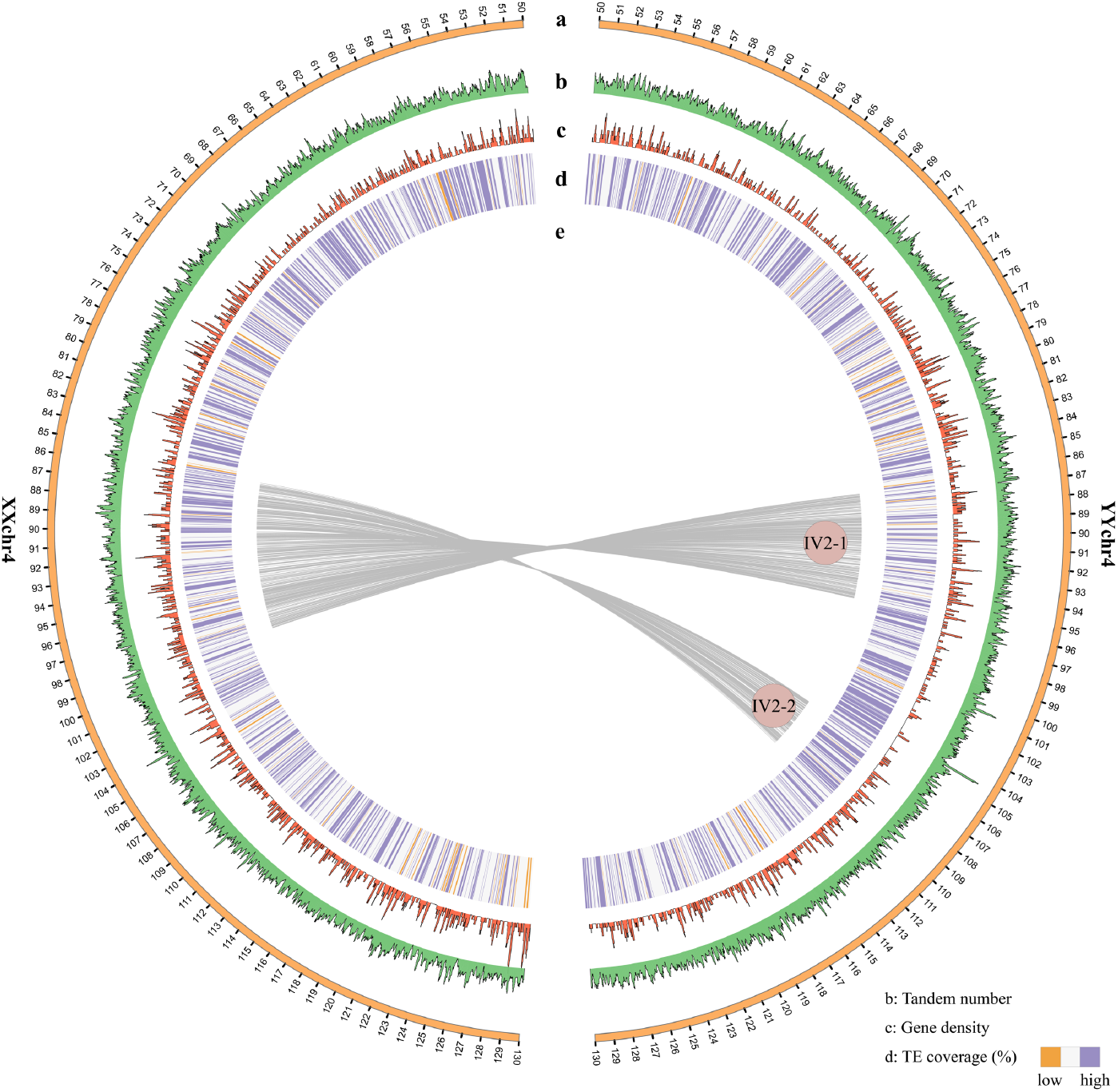
X and Y chromosome landscape. **(a)** Ideogram of the partial X and Y chromosome (in Mb scale). **(b)** Tandem number represented as the number of tandem repeats per 20 Kb. **(c)** Gene density in 20-Kb windows. **(d)** Percentage of coverage of transposable elements per 20-Kb. **(e)** Syntenic relationships of the sex determination region between the X and Y chromosome.

Subsequently, to estimate the evolutionary time since the suppression of recombination at the sex determination region based on homologous gene pairs, syntenic analysis was performed between the X and Y chromosome using MCscan (Figure 4A). A total of 84 and 35 syntenic gene pairs were identified within the IV2-1 and IV2-2, respectively, and they were used to calculate synonymous substitution rates (Ks) (remove abnormal value with high Ks value), which reflect the divergence time of the gene pairs between the X and Y chromosome (Table S14, Figure 4B). Additionally, 2,590 homologous genes within the non-inversion (non-IV) were also randomly selected to calculate Ks (Table S15). As shown in Figure 4B, the Ks value of IV2 clearly revealed two distinct evolutionary strata, corresponding perfectly with IV2-1 and IV2-2 (Figure 4A). The average Ks for the gene pairs located on IV2-2 was 0.0342, which was significantly higher than IV2-1 (average Ks = 0.0167). The heterogeneity of the Ks values of IV2 suggests that the recombination was suppressed at a different time for the sex determination region. Additionally, the Ks value of non-IV (average Ks = 0.0011) was significantly lower than IV2-1 and IV2-2, which reflects that recombination suppression might have occurred due to the two inversions. The divergence times for the gene pairs in IV2-1 and IV2-2 were on average 1.28 and 2.74 million years ago (Mya), respectively. Thus, we conclude that IV2-2 occurred firstly, followed by the expansion of the MSY, thus accounting for IV2-2 (~91%) containing more repetitive sequences than IV2-1 (87%) but less than MSY (92.32%) (Table S13).

**Figure 4.**
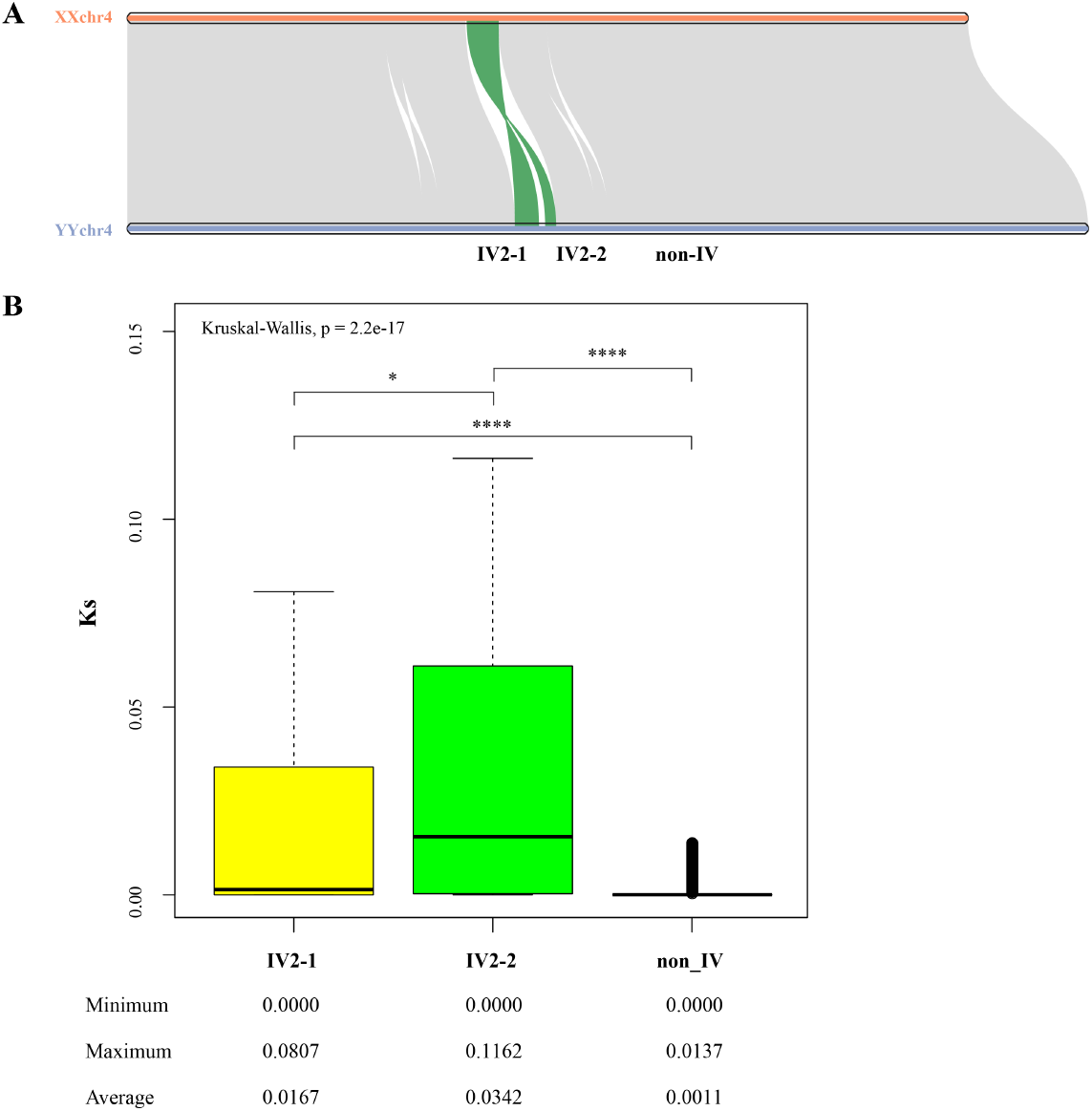
Synonymous divergence in the sex determination region of spinach. **(A)** Syntenic relationship of the X and Y chromosome. The green lines indicate the syntenic relationship of the sex determination region. **(B)** Boxplot of synonymous substitution rates for homologous gene pairs on IV2-1, IV2-2, and non-IV. A Krushal-Wallis test was used for comparison between the three regions (*P*<2.2e-17). IV: inversion; non-IV: non-inversion.

### Gene contents of the sex determination region of the X and Y chromosomes

Based on the genome annotation, 210 protein-coding genes were identified in the sex determination region of Y (Table S16) and 189 protein-coding genes were annotated in its X counterpart (Table S17), showing a lower gene content in the Y chromosome due to the MSY. The average gene density was nine genes per Mb in the Y chromosome, which was much lower than the average of 15 gene per Mb in the X counterpart, even in IV2-1 and IV2-2, which was still lower than the genome-wide average of 28 gene per Mb (Table 1 and Table 2). A total of 121 syntenic genes were shared by the X and Y chromosome, of which 84 and 35 genes were located at IV2-1 and VI2-2, respectively, whereas there were none at MSY. A total of 91 (43.3%) non-syntenic genes were found in the Y chromosome, of which 55 genes matched on autosomes, 23 genes were mapped on the X chromosome, and 13 genes did not share any homologous sequences with the X chromosome or autosome. Similarly, 67 non-syntenic genes (35.4%) were identified in the X chromosome, 39 genes were located at autosomes, 23 genes matched on the Y chromosome and 5 genes were without homologous genes (Figure S12). These non-syntenic genes in the Y chromosome were significantly related to photosynthesis and thylakoid and chlorophyll binding, while the non-syntenic genes in the X chromosome were enriched on telomere maintenance, DNA recombination, and DNA helicase activity, suggesting that the genes in the X and Y chromosome exerted different effects during spinach development (Figure S13 and Table S18).

**Table 2.** Summary of the genes in the sex determination region of the Y and X chromosomes

| Items | Y chromosome |  |  |  | X chromosome |  |  |
| --- | --- | --- | --- | --- | --- | --- | --- |
|  | MSY | IV2-1 | IV2-2 | Tot./Ave. | IV2-1 | IV2-2 | Tot./Ave. |
| Length (Mb) | 10.0 | 9.3 | 4.7 | 24.0 | 8.6 | 4.4 | 13.0 |
| Number of gene | 49 | 111 | 50 | 210 | 132 | 54 | 189* |
| Gene density (gene/Mb) | 5 | 12 | 11 | 9 | 15 | 12 | 15 |
| Syntenic gene number | 0 | 84 | 35 | 119 | 84 | 35 | 119 |
| Non-syntenic gene ratio (%) | 100 | 24.3 | 30.0 | 43.3 | 36.3 | 35.1 | 35.4 |
| Autosomal homologs (%) | 77.5 | 70.3 | 60.0 | 72.5 | 50.2 | 73.6 | 58.2 |

189\* included genes in IV2-1 (132) and IV2-2 (54) of the X chromosome and three genes between the IV2-1 and IV2-2. Tol., Total; Ave., Average.

The proportion of non-syntenic genes in IV2-1 was 24.3%, which was lower than IV2-2 (30.0%) in the Y chromosome, while the proportion appeared to be nonsignificant between IV2-1 (36.3%) and IV2-2 (35.1%) in the X chromosome (Table 2). Meanwhile, IV2-2 occurred prior to IV2-1 (Figure 4), and thus we hypothesized that the non-syntenic genes existing in the Y chromosome were more likely to have been gained (i.e., duplication from chromosomes) than lost from the X chromosome, which is supported by observations in papaya (Wang et al., 2012).

To search for candidate sex determinants, we mapped genomic reads from 20 male and 20 female individuals (Table S1) to the Y chromosome, which was used for identifying the male-specific genes (without female reads mapped). The results indicated that four male-specific genes (*YY_140950.1, YY_141010.1*, *YY_141140.1*, and *YY_141280.1*) were present in MSY. The partial genes within MSY exhibited male bias while genes flanked by MSY shared no substantial gender bias (Figure 5A). Of these four male-specific genes, only *YY_140950.1* and *YY_141140.1* were substantially male-specific expressed (TPM >1) among the three groups (Figure 5B and Table S20). Of the two fully male-specific genes (Figure 5B), *YY_140950.1* encodes a nucleic acid binding protein. The other gene, *YY_141140.1,* encodes a basic leucine zipper (bZIP) protein required for the positive regulation of flowering (Romera-Branchat et al., 2020). The presence of both gene sequences was restricted to the male in spinach (Figure S14A). Additionally, *YY_140950.1* and *YY_141140.1* were expressed in both the developing flowers and young leaves, whereas *YY_140950.1* was expressed faintly, demonstrating that the two genes were not specifically expressed (Figure S14B). The further investigation is necessary to confirm function of the two genes. Otherwise, based on the gene annotation, we focused on the genes related to floral organs (i.e., stamen and carpel) and hormones. A total of 12 genes were identified from the 210 genes in the Y chromosome, 50% of which were in MSY and six genes were detected as differentially expressed genes (DEGs) in at least one group (Table S20). Interestingly, a gene *YY_141510.1* encoding an R2R3 factor that is consistent with the sex determination gene *TDF1* in asparagus was detected, but the gene was located on IV2-2 and shared only one amino acid substitution of its homologous gene *XX_141830.1* on the X chromosome (Figure S15). Among the 210 genes, a total of 31 DEGs were found, 12 genes were detected only once, 19 genes were detected twice, which were mainly enriched on the MSY. Importantly, majority of the genes were up-regulated (male vs. female) (Figure 5 and Figure S16 and Table S20). Our analysis provides a basis to further identify and characterize sex determination genes in spinach.

**Figure 5.**
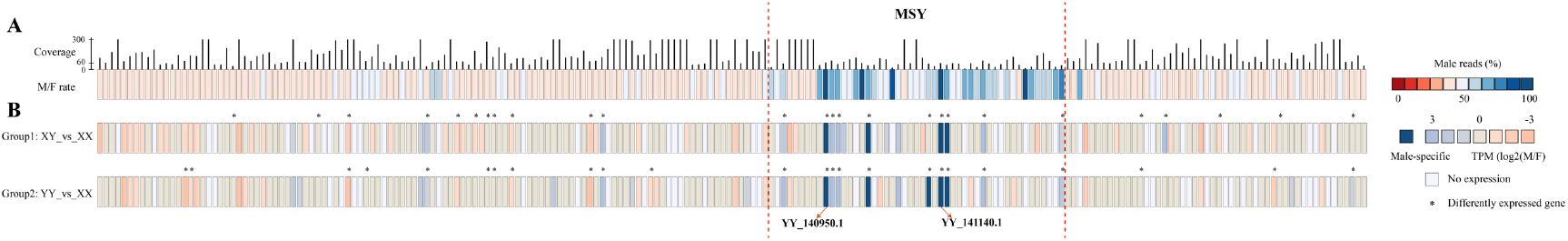
Characterization of candidate sex determining genes from 210 genes in the sex determination region. **(A)** Two hundred and ten genes with coverage of male genomic reads (black lines) and percentage of male genomic reads (color chart). Male-specific genes are shown in dark blue. The coverage at male-specific regions with no X chromosome counterpart is estimated at approximately 60×. **(B)** Expression bias between male and female individuals from the two groups in flowers. Complete male-specific expression is shown in dark blue. The gene with an asterisk indicates it’s a differently expressed gene. The individuals in group1 and group2 were from10S15. XX: female flower; XY: male flower with XY genotype; YY: male flower with YY genotype.

## Discussion

Dioecy, constituting a stable population with separate females and males, is a situation familiar to us, as the human population is divided into males and females according to inheritance of the Y chromosome (Henry et al., 2018). Investigating the sex determination mechanism of dioecious plants and animals is common. In the last few years, especially with the advent of the TGS technologies such as ONT, PacBio and 10X Genomics, are powerful techniques for assembling sex chromosomes and further identifying sex determination regions due to their ultra-long reads (Akagi et al., 2019; Jia et al., 2019). However, it is difficult to assemble the Y/W chromosome, as YY/WW viability is rare. Fortunately, the viable YY individual has been identified in spinach, thus offering the possibility to assembly the complete Y chromosome for uncovering the sex determination mechanism (Wadlington and Ming, 2018). In this study, female (XX) and male (YY) spinach individuals were used to assemble two genomes using ONT technology and a high-density genetic linkage map (Figure 1). Based on the genomes, a sex-determining region from 85.8–98.8 Mb on the X chromosome, corresponding 86.5–110.6 Mb on the Y chromosome, was identified, which was further validated by two independent approaches. The results demonstrated that the genomes generated herein exhibited better completeness and continuity than the spinach v1 genome (Xu et al., 2017) (Figure 2, Figure S10 and Figure S11). Additionally, a 10 Mb region from 95.8–105.9 Mb was defined as the male-specific region of the Y chromosome by sequence alignment and reported sex-linked/Y-specific markers (Figure 2). Multiple approaches were also used to demonstrate the authenticity of the MSY (Figure S9). Importantly, we first constructed the relative completeness of the X and Y chromosomes with lengths of about 184 Mb and 195 Mb, respectively, which was an available resource for investigating the sex determining mechanism in spinach.

Recombination suppression seems to be related to chromosomal rearrangements and in particular with inversion (Muyle et al., 2017; Nicolas et al., 2005; Wang et al., 2012; Yazdi and Ellegren, 2018). For example, two large inversion events define the two evolutionary strata in papaya (Wang et al., 2012). Even in animals, inversion events may also result in recombination suppression between the X and Y chromosome, such as in humans (Cotter et al., 2016), mole (Real et al., 2020), ostrich (Yazdi and Ellegren, 2018), and stickleback (Natri et al., 2019). Here, we did identify three large-scale inversions between the X and Y chromosome and further confirmed them using ONT long reads (Figure S6, Figure S7 and Figure S8). Interestingly, almost all reported sex-linked markers and fully sex-linked SNPs/InDels within the 59 individuals were substantially enriched on the inversion 2 (IV2) with a length of ~13 Mb, strongly suggesting that IV2 might be a sex determination region. IV2 was further divided into two parts named IV2-1 and IV2-2 based on the position of the MSY (Figure 2, Figure 3 and Figure 4). Specifically, IV2-1 was located at 86.5–95.8 Mb on the Y chromosome, corresponding to 90.3–98.8 Mb on the X chromosome, and IV2-2 was located at 105.9–110.6 Mb on the Y chromosome, with its X counterpart from 85.8 Mb to 90.1 Mb, both of which, importantly, were positioned on both sides of the MSY (Figure S7). Therefore, the recombination cessation of the MSY might be derived from IV2-1 and IV2-2 in spinach, which is the first case, to our knowledge, of two cross-inversion events suppress recombination, thus generating the MSY. Furthermore, the Ks value of IV2-2 was significantly higher than of IV2-1, suggesting that two evolutionary strata were required during the suppression of recombination (Figure 4 and Figure S17). The accumulation of repetitive sequences in the IV2-2 region was also higher than in the IV2-1 region, demonstrating that long periods of evolutionary time could result in a greater accumulation of repetitive sequences. A total of 92.32% of the MSY sequences are already repetitive, which is higher than in IV2-1 and IV2-2 (Table S13), and thus we propose that recombination stopped later than the evolution of the male-determining factor. Owing to two inversions (IV2-1 and IV2-2) flanked on both sides of the MSY, we hypothesize that a single inversion alone was not sufficient to maintain the stability of the MSY, and the stable MSY might have been generated when IV2-2 occurred (Figure S17). Actually, IV2-1 contain an approximately 1 Mb non-inversion region, whereas Ks values of the two regions (upstream and downstream of the 1 Mb) did not exhibit significant differences, and thus they were defined as the single inversion (IV2-1) in this study (Figure 2).

The MSY has increasingly expanded mainly due to the accumulation of high retrotransposons and other repetitive sequences during the evolution of the sex chromosome (Gschwend et al., 2012). In papaya, the hermaphrodite-specific region of the Y^h^ chromosome (HSY) pseudomolecule might have originated from the 157 pairs of intrachromosomal duplications (5.8% of the HSY sequence) (Wang et al., 2012). Here, a 10 Mb MSY was identified, harboring 92.32% repetitive sequences. How did the huge region generate during the evolution of spinach? Applying the criterion of at least 97% sequence identity of more than at least 5 Kb, we found 75 similar sequences (length of 791.1 kb) with 0–3% sequence divergence on each chromosome with fragment lengths ranging from 5 kb to 16.9 Kb (Table S21 and Figure S18). The Y chromosome encompassed the maximum number of similar sequences (N=22 or 28.9%), followed by YYchr2 (N=13 or 17.1%). Altogether, the intrachromosomal segmental duplications accounted for 1.74% of the MSY, which is lower than the interchromosomal duplications (3.91%). Additionally, the 49 genes in the MSY did not share synteny relationships between the Sp_XX_v1 and *C. quinoa* genomes, but matched on everywhere (67% on autosomes) in the Sp_YY_v1 genome (Figure S12, S19 and Table S16). Thus, we hypothesize that the MSY might be derived from duplication from autosomes. The assembly of *S. tetrandra* and *S. turkestanica,* both of which are wild-type individuals of spinach, will likely to uncover the evolutionary mechanism of MSY in the future.

In recent years, effective and efficient approaches have boosted the identification of sex-determining genes in dioecious plants. In kiwifruit, for example, two Y-specific genes, *SyGl* acting as the suppressor of feminization (SuF) and *FrBy* defining as maintenance of male (M) functions, were identified as sex-determinant candidate genes. *SyGl* encodes a type-C cytokinin response regulator (RR) that suppresses carpel development, while *FrBy* hinders tapetum degradation resulting in male sterility (Akagi et al., 2018; Akagi et al., 2019). An *RR* was also defined as a master regulator of sex determination in Salicaceae (Yang et al., 2020). In asparagus, defective in tapetal development and function 1 (*TDF1*), which promotes male function, and Suppressor of Female Function (*SOFF*), which suppresses female organogenesis were defined as sex determination genes, which supports the two-gene model. The Y-specific gene *TDF1* encodes an R2R3 MYB transcription factor, which exists as single nucleotide mutation induced by a premature stop codon, thus resulting in defective anthers (Harkess et al., 2017). Sex determination genes are always related to floral organ development (i.e., stamen or carpel), as mentioned above. Using this reasoning, we identified 12 genes related to pollen tube growth (Hoffmann et al., 2020; Li et al., 2017; Zhao et al., 2013) and male sterility (Torres et al., 2018) in the sex determination region of the Y chromosome (Table S20). Importantly, *YY_141510* encoded an R2R3 MYB transcription factor, which is consistent with the sex determination gene (M) in asparagus. However, the gene was located on IV2-2 and had a copy on the X chromosome (*XX_141830.1*), which shared an identical amino acid sequence, except for one SNP (Figure S15). Additionally, the two male-specific genes, *YY_140950.1* and *YY_141140.1,* identified based on genomic and transcriptome analysis, might not be direct regulators for determining sex in spinach, as *YY_140950.1* encodes a nucleic acid binding protein and *YY_141140.1* encodes a bZIP protein required for the positive regulation of flowering (Figure 5 and Figure S14). Indeed, the two genes were present in the male individuals of spinach, suggesting that these genes were conserved and exerted specific functions in the male individuals of spinach (Figure S14). Future studies should perform transgenic function experiments, virus-induced gene silencing (VIGS), or mutation (i.e., ethyl methanesulfonate (EMS), gamma irradiation and spontaneous mutants) experiments to further verify the function of candidate genes. Additionally, epigenetic could also regulate sex determination in animals and plants, such as miRNA (Akagi et al., 2016; Akagi et al., 2014) and methylation (Bräutigam et al., 2017; Kuroki et al., 2018), and thus we will conduct miRNA and methylation analysis of the genes within the sex-determining region in the future.

## Materials and methods

### Plant materials

A male individual from the inbred line 10S15 bearing some hermaphrodite flowers was self-pollinated to produce XX, XY, and YY individuals (Figure 6), in which XX and YY were used for ONT sequencing and *de novo* assembly, respectively. The identification of YY individuals was accomplished in our previous study (accepted). Eighty inbred lines (48 male and 32 female individuals) were used for resequencing and GWAS analysis in this study. Among them, 15 female and 16 male individuals were used for MSKs identification. Information on the 80 individuals is summarized in Table S1. All of the above plants were planted in the field at the Institute of Vegetables and Flowers (IVF) of the Chinese Academy of Agricultural Sciences (CAAS) in spring 2018. The floral morphology of each plant was visually inspected to determine the sexual phenotype.

**Figure 6.**
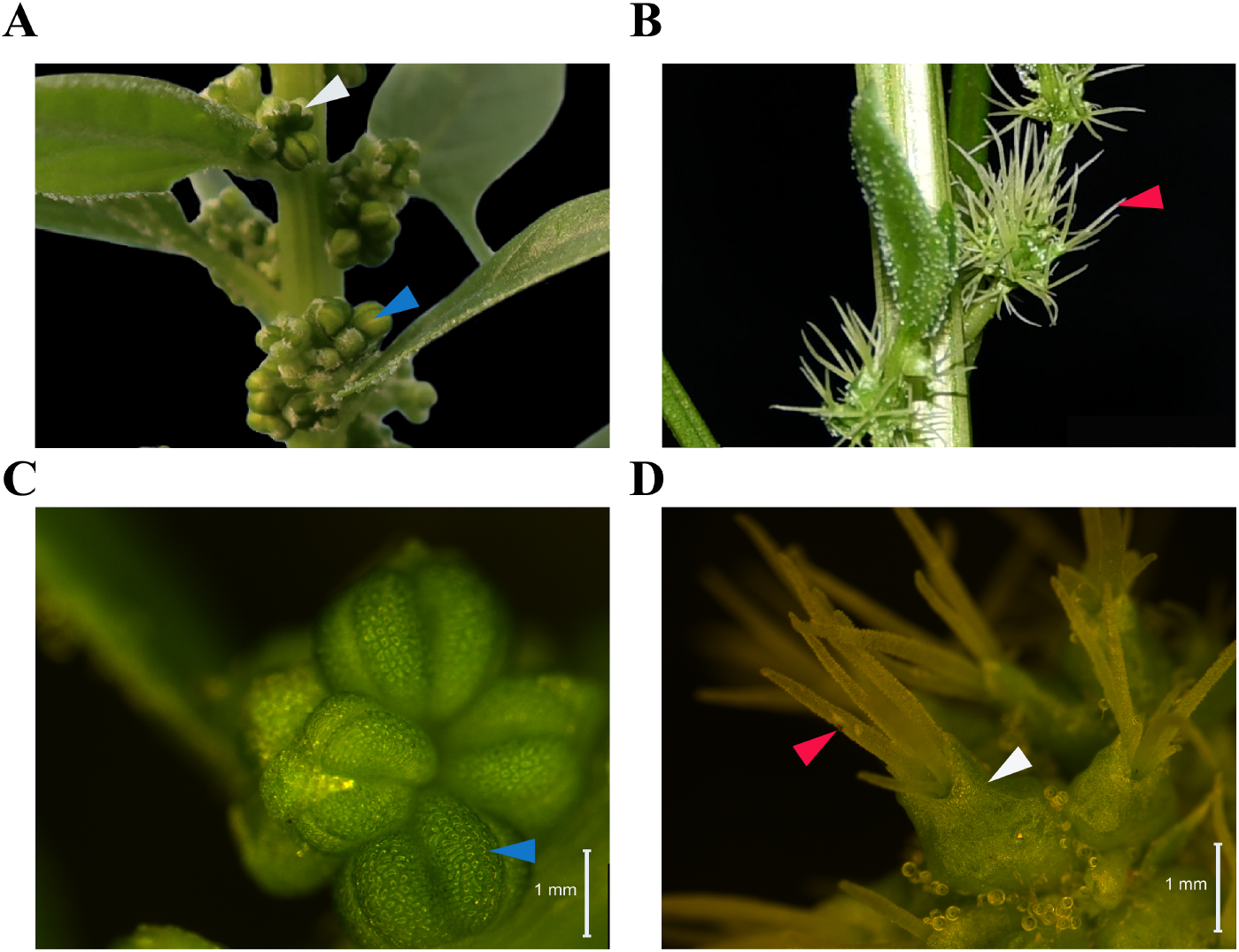
Characteristics of male (XY or YY) and female (XX) flower in spinach. **(A)** and **(C)** indicate male flower. **(B)** and **(D)** indicate female flower. White arrow, sepal; blue arrow, stamen; red arrow, stigma.

### Illumina library construction and sequencing

The fresh leaves from each individual were collected and frozen in liquid nitrogen prior to DNA extraction. High-quality genomic DNA was extracted as described by Murray and Thompson (1980) (Murray and Thompson, 1980). The DNA quality and concentration were assessed using electrophoresis on 1.0% agarose gels and an ND-2000 spectrophotometer (Thermo Fisher Scientific, Wilmington, DE, USA). The genomic DNA was used for Illumina and ONT library construction and then sequencing. Illumina library were sequenced using a HiSeq 2500 instrument (Illumina, San Diego, CA, USA) to generate 150 bp paired-end reads by BioMarker (Beijing, China). To correct errors in the Oxford Nanopore long reads, the Illumina genomic library from XX and YY used for ONT library construction was also sequenced on HiSeq 2500 (150-bp paired-end reads).

Early stage of inflorescence of female (XX), male (XY) and YY individuals with three biological replicates from inbred line 10S15 were collected on June 5, 2018. Total RNA was extracted using TRIzol reagent (Thermo Fisher Scientific Inc.) and purified by phenol/chloroform extraction. The cDNA was synthesized using a TranScript One-Step gDNA Removal and cDNA Synthesis Kit (TransGen Biotech, Beijing, China). The mRNA library was constructed and sequenced on a HiSeq 2500 (150-bp paired-end reads) by BerryGenomics (Beijing, China).

### Genome assembly of both female (XX) and male (YY) individuals

Nextdenovo (v2.2.0; https://github.com/Nextomics/NextDenovo) was used for assembling female (XX) and male (YY) individual genome. Specifically, the raw Oxford Nanopore long reads of females and males were assembled using Nextdenovo (parameters ‘read_cutoff = 1k, minimap2_options_raw = -x ava-ont -minlen 1000, random_round = 20’), three rounds of polishing were applied using Racon (v1.3.3; https://github.com/isovic/racon) with XX/YY ONT long reads, and two rounds of polishing were performed using NextPolish (v1.2.2; https://github.com/Nextomics) with the XX/YY Illumina reads (~50×). Additionally, NECAT (v0.01; https://github.com/xiaochuanle/NECAT) was also used to assemble the two individuals with default parameters to correct and extend the contigs. The assembled contigs were further polished using Medaka (v0.11.5; https://github.com/nanoporetech/medaka) and NextPolish. A high-density genetic linkage map (Qian et al., 2017) was applied to orient and order the contigs and adjacent contigs were separated by 100 ‘N’s. BUSCO was used to perform preliminary assessment of the assembly results using the embryophyta_odb10 database (Waterhouse et al., 2017). D-GENIES was used to perform synteny alignments between the female and male assembly (Cabanettes and Klopp, 2018). Hi-C reads from a male inbred line 12S4 were also used to assess the female and male genome. We aligned the Hi-C reads against the two assemblies and illustrated the whole-genome contact map using ALLHIC (https://github.com/tangerzhang/ALLHiC).

### Annotation of repetitive sequences

A *de novo* repetitive-element library was constructed using RepeatModeler (v.1.0.11) (http://repeatmasker.org/RepeatModeler.html). The repetitive elements of female (XX) and male (YY) genome were annotated using RepeatMasker (v.4.0.7) (http://repeatmasker.org/) (Zhang et al., 2012). Additionally, tandem repeats were analyzed using Tandem Repeats Finder software (TRF) (v4.09) (Benson, 1999).

### Protein-coding gene prediction and functional annotation

The female (XX) and male (YY) genomes were used for gene prediction using MAKER (v2.31.6) (Cantarel et al., 2008), which combines evidence from *ab initio*, transcript mapping and protein homology-based predictions. SNAP (v2006-07-28) (Korf, 2004), AUGUSTUS (v3.3.2) (Stanke et al., 2006), and GeneMark (v4.57_lic) (Besemer and Borodovsky, 2005) were used for *ab initio* gene predictions. For transcript mapping, 18 RNA-Seq data from NCBI, nine RNA-seq data from female (XX), male (XY), and YY individuals from 10S15 were filtered using fastp (v0.20.0) (https://github.com/OpenGene/fastp#get-fastp) (Table S6). The RNA-Seq data were then aligned to the XX/YY genome using STAR (v1.5.2) (Dobin et al., 2012) and then assembled using Stringtie v2.1.2 (Kovaka et al., 2019) and gffread (https://github.com/gpertea/gffread). Protein sequences from sugar beet, *Arabidopsis*, and Swiss-Prot were used for protein homology-based prediction. The predicted genes without start codons, stop codons, or with a length less than 50 bp were filtered. To annotate the predicted genes, their protein sequences were mapped to the *Arabidopsis* protein, Swiss-Prot, and TrEMBL (http://www.uniprot.org/) databases using BLASTP (‘-evalue 1e-5’), as well as the InterPro database using InterProScan (v5.10-50.0) (Jones et al., 2014). Functional descriptions and GO annotations were performed using AHRD (v3.3; https://github.com/groupschoof/AHRD).

We also performed noncoding RNA annotation. The tRNA annotation was conducted using tRNAscan-SE (v1.3.1) with the default parameters (Lowe and Eddy, 1997). The rRNA annotation was performed using RNAmmer (v1.2, ‘-S euk -m lsu,ssu,tsu -multi’) (Lagesen et al., 2007). Others RNAs were annotated by infernal (v1.1.2) (Nawrocki and Eddy, 2013) using the Rfam 14.0 database (Griffiths-Jones et al., 2005).

### Phylogenetic inference

Single copy genes were identified using OrthoFinder (v2.2.7) (Emms and Kelly, 2015) with an all – versus-all BLASTP comparison of protein sequences from spinach and seven other plant species, namely, *A. hypochondriacus*, *A. thaliana*, *C. quinoa*, *V. vinifera*, *O. sativa*, *B. vulgaris* and *S. lycopersicum*. The protein sequences of the single copy gene families within the species were aligned using MAFFT (v7.158b) (Katoh et al., 2005), and conserved aligned regions were extracted using Gblock (v0.91b) with parameters ‘-t=p, -b4=5, -b5=h’. Finally, the concatenated protein alignments were used to construct a phylogenetic tree using the maximum parsimony method in PhyML (v3.0) (Guindon and Gascuel, 2003).

### Analysis of MSY

To define the MSY, a comparison of the X and Y chromosome was performed using minimap2 (v 2.15-r915-dirty) (https://github.com/lh3/minimap2), and a putative MSY of 10 Mb, was identified. Five independent approaches were used to validate the accuracy of the MSY as follows: (1) published sex-linked and male-specific markers (Arumuganathan and Earle, 1991; Groben and Wricke, 1998; Kudoh et al., 2017; Okazaki et al., 2019; Wadlington and Ming, 2018) were aligned to the X and Y chromosome, respectively using BLASTN (‘-evalue 1e-10’). The sex-linked markers and male-specific markers were then used to determine the MSY; (2) 49 samples (21 female, 37 male, and one YY individuals) of resequencing data (Table S1) were aligned to the Y chromosome using BWA (v0.7.17) with the default parameters, and fully sex-linked SNPs and InDels, heterozygous X^a^Y^b^ in males but homozygous X^a^X^a^, Y^b^Y^b^ in females, and YY individuals were identified using SAMtools (v1.6-3) (Li et al., 2009) and BCFtools (v1.6) (Danecek et al., 2011); (3) based on the alignment results of step (2), the average depth of coverage of 20 females and 20 males was extracted for each 10 Kb using SAMtools (v1.6-3) (Table S1). A comparison of the relative depth of coverage between male and female individuals revealed the MSY; (4) to verify the MSY with low recombination, SLAF markers from the high-density genetic map were mapped to the Y chromosome using BLASTN (‘-evalue 1e-10’); (5) a female spinach reference genome (Dohm et al., 2013) was mapped to the Y chromosome to validate the accuracy of the MSY using minimap2.

### Identification of sex-related contigs using reference free k-mer analysis

Fifteen female and 16 male individuals from the inbred lines were used to identify sex-related sequences through comparison of the specific lengths of sequence numbers between the female and male sequence pools. In brief, all 31-bp k-mers were selected from all genomic reads for females and males using Jellyfish (v2.3.0) (Marcais and Kingsford, 2011). To identify the male-specific k-mers (MSKs) that were completely absent in the female reads, k-mer counts were compared between the male and female reads using a custom script. The most robust set of MSKs was generated by selecting the MSK that met a total count ≥20 and ≤ 200, which could avoid sequencing error and repetitive sequences.

One hundred and fifty-bp paired-end (PE) reads that included at least one MSK were extracted using Cookiecutter (v1.0.0; https://github.com/ad3002/Cookiecutter) (Starostina et al., 2015). The PE reads containing MSKs were aligned to the Sp_YY_v1 and spinach v1 genomes using Burrows-Wheeler Aligner (BWA) (v0.7.17) (Li and Durbin, 2009) with default parameters. The number of mapped reads per 20-kb bin was counted as described by (Akagi et al., 2018). *De novo* assembly was conducted using SOAPdenovo (v2.04) (Luo et al., 2012), and the contigs that met a minimum length threshold of 200 were retained. To identify sex-linked and male-specific contigs, we map the 31 genomic reads (15 female and 16 male individuals) against these contigs using BWA (v0.7.17) with the default parameters. Informative SNPs were identified using SAMtools (v1.6-3) (Li et al., 2009) and BCFtools (v1.6) (Danecek et al., 2011) using the following parameters: mpileup -q 20 -Q 20 -C50 -Ou -f ref.fa | bcftools call -cv. The informative SNPs were supported by at least three distinct reads. Based on the alignment above, the contigs were further classified into two categories: Y-specific contigs and sex-linked contigs. Briefly, the contigs with no female reads mapped were considered as Y-specific. The contigs bearing significant SNPs that co-segregated with > 90% individuals, were judged to sex-linked contigs. Additionally, to anchor the sex-linked contigs to Sp_YY_v1, we sought to map these contigs against the Sp_YY_v1 genome using BLASTN (v2.8.1), allowing up to 10% mismatches in a 100-bp bin.

### Genome-Wide Association Studies

Eighty samples (48 male and 32 female individuals) of resequencing data were used for GWAS analysis (Table S1). Firstly, the low-quality reads and adapters of 80 individuals were filtered using fastp (v0.20.0; parameters ‘-q 20’; https://github.com/OpenGene/fastp#get-fastp). Clean reads were then aligned to the spinach v1 genome (Xu et al., 2017), and a set of SNPs was identified using PopSeq2Geno (Cheng et al., 2016). The identified SNPs were supported by a minor allele frequency (MAF) >= 0.05 and missing genotype < 5%. Next, GWAS was performed using the compressed mixed linear model (CMLM) within the GAPIT package of R (v3.4.2). The three principal components (PCs) generated from principal component analysis (PCA) were used to estimate the population structure. The significance threshold was set to *P* = 0.05 using the false discovery rate (FDR) correction (Benjamini and Hochbery, 1995). Quantile-quantile (QQ) plots and Manhattan plots were plotted using the qqman package of R (v3.4.2) (Turner, 2014).

### Inversion analysis

Based on the alignment results of the X and Y chromosome, three large-scale inversions were detected between the X and Y chromosome. To validate the three inversions, the XX and YY ONT reads were aligned to the X and Y chromosome using minimap2. For an inversion on the Y chromosome, more than two YY ONT reads were perfectly mapped to the border of the inversion, and the outer border shared the homolog region with its X counterpart while the inner border shared the inverted region with its X counterpart, suggesting that an inversion does exist between the X and Y chromosome.

### Similarity, synteny analysis, and divergence time estimation

Similarity sequence analysis of the MSY and whole male (YY) genome was conducted using NUCmer (v3.1, parameters ‘-I 97 -L 5000’) (http://mummer.sourceforge.net/manual/). The similar sequence met the minimum length of 5,000 bp and 97% identity.

The synteny blocks between the X and Y chromosome were identified and visualized using MCscan (Python-version) (parameter ‘--minspan=30’, https://github.com/tanghaibao/jcvi/wiki/MCscan-(Python-version)). Images of repetitive sequences, gene density, and synteny blocks of the X and Y chromosome were drawn using Circos (Krzywinski et al., 2009). The yn00 program of PAML v4.9j (Yang, 2007) was used to estimate synonymous divergence (dS) between the X/Y gene pair. A boxplot of dS values was generated with R software (v3.6.1). Divergence times were estimated as T = K/2r and the r = 6.5 × 10^−9^ as recommended by (Okazaki et al., 2019).

### Identification of candidate sex determining genes

The GO enrichment analysis was performed using clusterProfiler (Yu et al., 2012). To estimate the male-specific genes at the genomic level, the reads counts per gene were generated from the aligned BAM files of the 20 female and 20 male individuals mentioned above using a custom R script. The genes were defined as a male-specific when the reads coverage ratio of male/female > 95%. Additionally, mRNA-seq reads from three female (XX), three male (XY) and three YY individuals from inbred line 10S15 were used to identify the genes substantially expressed in developing flowers. These mRNA-seq reads were aligned to the Y chromosome using HISAT2 (v4.8.2) with default parameters (Kim et al., 2015). The read count per gene was generated using featureCounts (v2.0.1) (Yang et al., 2014), and the read count per gene was converted to transcripts per kilobase million (TPM) using a custom python script. The mRNA-seq data were further divided into two groups. Three female (XX) and three male (XY) individuals were defined as group1; and the three female (XX) and three YY individuals were defined as group2. Differential expression between male (XY)/YY and female individuals was analyzed in R (v3.4.2) using the R package DESeq (v1.14) (Anders and Huber, 2010). *P* < 0.05 and Fold Change (FC) > 2 were used to identify DEGs. The genes were defined as male-specific expressed when the average TPM = 0 and average TPM > 1 in the female and male, respectively. Veen diagram of the DEGs in three groups using BMKCloud (http://www.biocloud.net/).

The primers used in the study were designed using online software Primer 3 (v0.4.0; https://bioinfo.ut.ee/primer3-0.4.0/). The PCR was performed in a total reaction volume of 10 μL containing 5 μL 2×Taq Master Mix (CoWin Biosciences, China), 0.25 μL forward and reverse primer, respectively, 3.6 μL ddH_2_O and 1 μL DNA/cDNA. The reaction was performed on a Veriti 96 Well Thermal Cycler (Applied Biosystems, Foster City, CA, US) under the following conditions: 5 min at 94°C; followed by 35 cycles of 30 s at 94°C, 30 s at 60°C, and 40 s at 72°C. The result was assessed using electrophoresis on 1.5% agarose gels.

## Data availability

Sequence data presented in this article could be download from the ####. Accession numbers are listed in ###. The genome assemblies are available through Phytozome ###.

## Acknowledgements

This work was performed at the Key Laboratory of Biology and Genetic Improvemet of Horticultural Crops, Ministry of Agriculture, Beijing, China, and was supported by the Natural Science Foundation of China (31872102), the Chinese Academy of Agricultural Sciences Innovation Project (CAAS-ASTIP-IVFCAAS), Beijing Scientific Program of Municipal Commission of Science and Technology (Z171100001517014), Central Public-interest Scientific Institution Basal Research Fund(IVF-BRF2018004), and the National Key Research and Development Program of China (2018YFD0100805).

## Authors’ contributions

WQ designed the study. HS conducted the experiments. HS and ZL analyzed the data. HS wrote the manuscript. WQ and XW revised the manuscript. WQ, ZL, HZ and ZX prepared the samples. FC helped analyzed data.

## Supplementary information

**Figure S1** An example of the correction and extension of contigs using two software programs and a linkage map. **(A)** Dot-plot alignment of some contigs from the XX genome assembly by Nextdenovo and NECAT. ND_XX contigs: the female contigs assembled by Nextdenovo; NC_XX contigs: the female contigs assembled by NECAT. **(B)** The position of SLAF markers on the contig199 and the extended contig **(C)**. LG4: linkage group 4.

**Figure S2** Alignment of the female **(A)** and male **(B)** assembled genome with the SLAF markers linkage genetic map. LG: linkage group.

**Figure S3** Dot-plot alignment of the female (Sp_XX_v1) with male (Sp_YY_v1) genome.

**Figure S4** Whole-genome contacts of the Hi-C data of male **(A)** and female **(B)** genomes.

**Figure S5** Genome evolution and comparative analysis of the spinach genome. **(A)** Phylogenetic analysis of spinach and other plants. **(B)** Syntenic relationships between male (YY) spinach, sugar beet and quinoa genomes.

**Figure S6** Illustration of the first inversion (IV1) between the X and Y chromosome using XX and YY ONT reads, respectively. The numbers shown on the bottom (top) of the X (Y) chromosome indicate the breakpoint of the inversion.

**Figure S7** Illustration of the second inversion (IV2) between the X and Y chromosome using XX and YY ONT reads, respectively. The numbers shown on the bottom (top) of the X (Y) chromosome indicate the breakpoint of the inversion.

**Figure S8** Illustration of the third inversion (IV3) between the X and Y chromosome using XX and YY ONT reads, respectively. The numbers shown on the bottom (top) of the X (Y) chromosome indicate the breakpoint of the inversion.

**Figure S9** Validation of the male-specific region (MSY) of Y in spinach. **(A)** Dot plot of a female genome assembled by Dohm et al. (2014) and the Y chromosome. **(B)** Sex mapped reads count ratio (log(F/M)) for 20 female and 20 male individuals are shown by black dots. The mapped reads per 10-kb bin was counted. The MSY is shaded. **(C)** Enrichment of fully sex-linked markers with 59 samples (21 female, 37 male, and one YY individuals) on Y chromosome. The number of sex-linked markers were counted for each 1 Mb. IV2 indicates the second inversion between X and Y chromosome. **(D)** The position of SLAF markers on Y chromosome. LG4: linkage group 4.

**Figure S10** Validation of the sex-determining region using GWAS and reference free k-mer. **(A)** Manhattan plot based on the results of GWAS. The y-axis represents the strength association (-log_10_(*P* value)) for each SNP. The red line indicates the threshold value (α < 0.05). **(B)** Quantile-quantile (QQ) plot derived from genome-wide association study. **(C)** Summary of mainly SNPs significantly associated with sex. SSC: Super_scaffold, Homo: Homozygous, Heter: Heterozygosis. **(D)** Mapping of reads including male-specific k-mers (MSKs) to the spinach v1 genome. **(E)** Enrichment of sex-linked contigs on spinach v1 genome. The box indicates chromosomes/scaffolds, and the name is shown on the left. The lines within the box represent the sex-linked contigs and the sex-linked regions have been highlighted with red lines.

**Figure S11** Validation of the sex-related sequences identified by GWAS and k-mers approach using the male (YY) genome. **(A)** Mapping of reads including male-specific k-mers (MSKs) to the male (YY) genome. **(B)** Enrichment of sex-linked/Y-specific contigs on the Y chromosome. The 93.4% Y-specific contigs and 70.9% sex-linked contigs are enriched on IV2 and MSY. MSY: male-specific region of Y chromosome. IV2: the second inversion between the X and Y chromosome. **(C)** The position of five main sex-associated regions on the Y chromosome. Chr4-sex and Chr3-sex indicates two regions associated with sex identified in figure S10. **(D)** Alignment of the chromosome 3 from the spinach v1 genome with the SLAF markers linkage genetic map. LG: linkage group.

**Figure S12** Distribution of the protein-coding genes in the sex determination region of the **(A)** Y and **(B)** X chromosomes. The syntenic genes are identified using the MCscan. The orthologous genes are identified using blast with the ‘–evalue 1e-10’.

**Figure S13** GO enrichment of non-syntenic genes in sex-determining region of **(A)** the Y and **(B)** X chromosomes.

**Figure S14** Characterization of the *YY_140950* and *YY_141140* genes. **(A)** Complete male-specific conservation of *YY_140950.1* and *YY_141140.1* in the genomes of *S. oleracea*. F: female; M: male. **(B)** Expression pattern of *YY_140950.1* and *YY_141140.1* in the flower and leaf of a male individual from inbred line 10S15. The *Spo01815* is considered as an actin gene from the spinach v1 genome.

**Figure S15** Amino acid alignment of *YY_141510* and *XX_141830*. Red box indicates amino acid substitution.

**Figure S16** Characterization of the differentially expressed genes (DEGs) identified from the sex determination region of Y chromosome. **(A)** Veen diagram of the DEGs in three groups. **(B)** Distribution of the DEGs in the IV2-1, IV2-2 and MSY.

**Figure S17** Evolution of the male-specific region of Y chromosome in spinach. **(A)** The Proto-X and Proto-Y chromosome in spinach. **(B)** The IV2-2 was occurred firstly between the proto-X and proto-Y chromosome, resulting recombination suppression, repetitive sequences accumulated and generation of the MSY **(C)**. **(D)** Subsequently, the IV2-1 was occurred between X and Y chromosome, which could maintain the stableness of the MSY, thus generating larger region of the MSY **(E)**.

**Figure S18** Segmental duplications of the MSY and whole genome. Each line within the circus indicates similar fragment with ≥5 kb length and ≥97% identity.

**Figure S19** Syntenic relationships between the MSY, Sp_XX_v1 and *C. quinoa* genomes. Corresponding syntenic regions are drawn. The bar with blue, orange and light green indicate IV2-1, MSY and IV2-2, respectively.

**Table S1** Statistics of the samples used in the study.

**Table S2** Assembly statistics of the female and male individuals.

**Table S3** The statistics of Illumina reads mapped to female and male genomes.

**Table S4** Summary of the BUSCO groups searched in female and male genome.

**Table S5** The statistics of repetitive sequences in the femal and male genome assembly.

**Table S6** The RNA-seq data used for annotation in the study.

**Table S7** Summary of the functional annotation in female and male genome.

**Table S8** List of the plant genome sequences used in the comparative genomic analysis.

**Table S9** Results of GWAS for sex in spinach.

**Table S10** The position of 304 male-specific contigs on the Y chromosome.

**Table S11** The position of 1362 sex-linked contigs on the Y chromosome.

**Table S12** Transposable elements in the sex-determining region.

**Table S13** Transposable elements in the IV2-1, IV2-2, and MSY.

**Table S14** Estimated synonymous nucleotide divergence of sex determination gene pairs between X and Y chromosome.

**Table S15** Estimated synonymous nucleotide divergence of non-inversion region gene pairs between X and Y chromosome.

**Table S16** Gene annotation of the sex determination region of the Y chromosome.

**Table S17** Gene annotation of the sex determination region of the X chromosome.

**Table S18** GO enrichment analysis of non-syntenic genes within the sex determination region of X and Y chromosomes.

**Table S19** Expression patterns of the 210 genes identified in the sex determination on Y chromosome between female (flower) and male (flower).

**Table S20** Summary of candidate genes identified in sex determination region on Y chromosome.

**Table S21** Segmental duplications on the MSY of Y chromosome.

## References

Akagi, T., Henry, I.M., Kawai, T., Comai, L. and Tao, R.J.T.P.C. (2016) Epigenetic regulation of the sex determination gene MeGI in polyploid persimmon. 28, 2905–2915.

Akagi, T., Henry, I.M., Ohtani, H., Morimoto, T., Beppu, K., Kataoka, I. and Tao, R. (2018) A Y-Encoded Suppressor of Feminization Arose via Lineage-Specific Duplication of a Cytokinin Response Regulator in Kiwifruit. Plant Cell 30, 780–795.

Akagi, T., Henry, I.M., Tao, R. and Comai, L. (2014) A Y-chromosome–encoded small RNA acts as a sex determinant in persimmons. Science 346, 646–650.

Akagi, T., Pilkington, S.M., Varkonyi-Gasic, E., Henry, I.M., Sugano, S.S., Sonoda, M., Firl, A., McNeilage, M.A., Douglas, M.J., Wang, T., Rebstock, R., Voogd, C., Datson, P., Allan, A.C., Beppu, K., Kataoka, I. and Tao, R. (2019) Two Y-chromosome-encoded genes determine sex in kiwifruit. Nat Plants 5, 801–809.

Akamatus, T., Suzuki, T. and Uchimiya, H. (1998) Determination of male or female of spinach by using DNA marker., Japan: Sakata no tane KK. Anders, S. and Huber, W. (2010) Differential expression analysis for sequence count data. Genome Biol 11, R106.

Arumuganathan, K. and Earle, E.D. (1991) Nuclear DNA Content of Some Important Plant Species. Plant Molecular Biology Reporter 9, 208–218.

Benjamini, Y. and Hochbery, Y. (1995) Controlling The False Discovery Rate: A Practical and Powerful Approach to Multiple Testing. Methodological 57, 289–300.

Benson, G. (1999) Tandem repeats finder: a program to analyze DNA sequences. Nucleic Acids Research 27, 573–580.

Besemer, J. and Borodovsky, M. (2005) GeneMark: web software for gene finding in prokaryotes, eukaryotes and viruses. Nucleic Acids Research 33, W451–W454.

Bräutigam, K., Soolanayakanahally, R., Champigny, M., Mansfield, S., Douglas, C., Campbell, M.M. and Cronk, Q.J.S.r. (2017) Sexual epigenetics: gender-specific methylation of a gene in the sex determining region of Populus balsamifera. 7, 1–8.

Cabanettes, F. and Klopp, C.J.P. (2018) D-GENIES: dot plot large genomes in an interactive, efficient and simple way. 6, e4958.

Cai, X., Xu, C., Wang, X., Wang, S., Zhang, Z., Fei, Z. and Wang, Q. (2018) Construction of genetic linkage map using genotyping-by-sequencing and identification of QTLs associated with leaf color in spinach. Euphytica 214.

Cantarel, B.L., Korf, I., Robb, S.M., Parra, G., Ross, E., Moore, B., Holt, C., Sanchez Alvarado, A. and Yandell, M. (2008) MAKER: an easy-to-use annotation pipeline designed for emerging model organism genomes. Genome Res 18, 188–196.

Charlesworth, B. and Charlesworth, D. (2018) A model for the evolution of dioecy and gynodioecy. American Society of Naturalists 112, 975–997.

Charlesworth, D. (2013) Plant sex chromosome evolution. Journal of Experimental Botany 64, 405–420.

Charlesworth, D. (2019) Young sex chromosomes in plants and animals. New Phytol 224, 1095–1107.

Cheng, F., Sun, R., Hou, X., Zheng, H., Zhang, F., Zhang, Y., Liu, B., Liang, J., Zhuang, M., Liu, Y., Liu, D., Wang, X., Li, P., Liu, Y., Lin, K., Bucher, J., Zhang, N., Wang, Y., Wang, H., Deng, J., Liao, Y., Wei, K., Zhang, X., Fu, L., Hu, Y., Liu, J., Cai, C., Zhang, S., Zhang, S., Li, F., Zhang, H., Zhang, J., Guo, N., Liu, Z., Liu, J., Sun, C., Ma, Y., Zhang, H., Cui, Y., Freeling, M.R., Borm, T., Bonnema, G., Wu, J. and Wang, X. (2016) Subgenome parallel selection is associated with morphotype diversification and convergent crop domestication in Brassica rapa and Brassica oleracea. Nature Genetic 48, 1218–1224.

Cotter, D.J., Brotman, S.M. and Sayres, M.A.W.J.G. (2016) Genetic diversity on the human X chromosome does not support a strict pseudoautosomal boundary. 203, 485–492.

Danecek, P., Auton, A., Abecasis, G., Albers, C.A., Banks, E., DePristo, M.A., Handsaker, R.E., Lunter, G., Marth, G.T., Sherry, S.T., McVean, G., Durbin, R. and Genomes Project Analysis, G. (2011) The variant call format and VCFtools. Bioinformatics 27, 2156–2158.

Deng, C., Qin, R., Gao, J., Cao, Y., Li, S., Gao, W. and Lu, L. (2013) Identification of sex chromosome of spinach by physical mapping of 45s rDNAs by FISH. Caryologia 65, 322–327.

Dobin, A., Davis, C.A., Schlesinger, F., Drenkow, J., Zaleski, C., Jha, S., Batut, P., Chaisson, M. and Gingeras, T.R. (2012) STAR: ultrafast universal RNA-seq aligner. Bioinformatics 29, 15–21.

Dohm, J.C., Minoche, A.E., Holtgräwe, D., Capella-Gutiérrez, S., Zakrzewski, F., Tafer, H., Rupp, O., Sörensen, T.R., Stracke, R., Reinhardt, R., Goesmann, A., Kraft, T., Schulz, B., Stadler, P.F., Schmidt, T., Gabaldón, T., Lehrach, H., Weisshaar, B. and Himmelbauer, H. (2013) The genome of the recently domesticated crop plant sugar beet (Beta vulgaris). Nature 505, 546–549.

Emms, D.M. and Kelly, S.J.G.b. (2015) OrthoFinder: solving fundamental biases in whole genome comparisons dramatically improves orthogroup inference accuracy. 16, 157.

Griffiths-Jones, S., Moxon, S., Marshall, M., Khanna, A., Eddy, S.R. and Bateman, A.J.N.a.r. (2005) Rfam: annotating non-coding RNAs in complete genomes. 33, D121–D124.

Groben, R. and Wricke, G. (1998) Occurrence of microsatellites in spinach sequences from computer databases and development of polymorphic SSR markers. Plant Breeding 117, 271–274.

Gschwend, A.R., Weingartner, L.A., Moore, R.C. and Ming, R.J.C.r. (2012) The sex-specific region of sex chromosomes in animals and plants. 20, 57–69.

Guindon, S. and Gascuel, O.J.S.b. (2003) A simple, fast, and accurate algorithm to estimate large phylogenies by maximum likelihood. 52, 696–704.

Harkess, A., Zhou, J., Xu, C., Bowers, J.E., Van der Hulst, R., Ayyampalayam, S., Mercati, F., Riccardi, P., McKain, M.R., Kakrana, A., Tang, H., Ray, J., Groenendijk, J., Arikit, S., Mathioni, S.M., Nakano, M., Shan, H., Telgmann-Rauber, A., Kanno, A., Yue, Z., Chen, H., Li, W., Chen, Y., Xu, X., Zhang, Y., Luo, S., Chen, H., Gao, J., Mao, Z., Pires, J.C., Luo, M., Kudrna, D., Wing, R.A., Meyers, B.C., Yi, K., Kong, H., Lavrijsen, P., Sunseri, F., Falavigna, A., Ye, Y., Leebens-Mack, J.H. and Chen, G. (2017) The asparagus genome sheds light on the origin and evolution of a young Y chromosome. Nature Communications 8, 1279.

Henry, I.M., Akagi, T., Tao, R. and Comai, L. (2018) One hundred ways to invent the sexes theoretical and observed paths to dioecy in plants. Annual Reviews 69, 553–575.

Hoffmann, R.D., Portes, M.T., Olsen, L.I., Damineli, D.S.C., Hayashi, M., Nunes, C.O., Pedersen, J.T., Lima, P.T., Campos, C., Feijo, J.A. and Palmgren, M. (2020) Plasma membrane H(+)-ATPases sustain pollen tube growth and fertilization. Nat Commun 11, 2395.

Iizuka, M. and Janick, J. (1962) Cytogenetic analysis of sex determination in Spinacia oleracea. Genetics 47, 1225–1241.

Jia, H.-M., Jia, H.-J., Cai, Q.-L., Wang, Y., Zhao, H.-B., Yang, W.-F., Wang, G.-Y., Li, Y.-H., Zhan, D.-L., Shen, Y.-T., Niu, Q.-F., Chang, L., Qiu, J., Zhao, L., Xie, H.-B., Fu, W.-Y., Jin, J., Li, X.-W., Jiao, Y., Zhou, C.-C., Tu, T., Chai, C.-Y., Gao, J.-L., Fan, L.-J., van de Weg, E., Wang, J.-Y. and Gao, Z.-S. (2019) The red bayberry genome and genetic basis of sex determination. Plant Biotechnology Journal 17, 397–409.

Jones, P., Binns, D., Chang, H.-Y., Fraser, M., Li, W., McAnulla, C., McWilliam, H., Maslen, J., Mitchell, A., Nuka, G., Pesseat, S., Quinn, A.F., Sangrador-Vegas, A., Scheremetjew, M., Yong, S.-Y., Lopez, R. and Hunter, S. (2014) InterProScan 5: genome-scale protein function classification. Bioinformatics 30, 1236–1240.

Katoh, K., Kuma, K.-i., Toh, H. and Miyata, T.J.N.a.r. (2005) MAFFT version 5: improvement in accuracy of multiple sequence alignment. 33, 511–518.

Kim, D., Langmead, B. and Salzberg, S.L.J.N.M. (2015) HISAT: a fast spliced aligner with low memory requirements. Nature Methods 12, 357–360.

Korf, I. (2004) Gene finding in novel genomes. BMC Bioinformatics 5, 59.

Kovaka, S., Zimin, A.V., Pertea, G.M., Razaghi, R. and Pertea, M.J.G.b. (2019) Transcriptome assembly from long-read RNA-seq alignments with StringTie2. Genome Biology 20.

Krzywinski, M., Schein, J., Birol, I., Connors, J., Gascoyne, R., Horsman, D., Jones, S.J. and Marra, M.A. (2009) Circos: an information aesthetic for comparative genomics. Genome Res 19, 1639–1645.

Kudoh, T., Takahashi, M., Osabe, T., Toyoda, A., Hirakawa, H., Suzuki, Y., Ohmido, N. and Onodera, Y. (2017) Molecular insights into the non-recombining nature of the spinach male-determining region. Molecular Genetics and Genomics 293, 557–568.

Kuroki, S., Tachibana, M.J.M. and endocrinology, c. (2018) Epigenetic regulation of mammalian sex determination. 468, 31–38.

Lagesen, K., Hallin, P., Rødland, E.A., Stærfeldt, H.-H., Rognes, T. and Ussery, D.W.J.N.a.r. (2007) RNAmmer: consistent and rapid annotation of ribosomal RNA genes. 35, 3100–3108.

Li, D.D., Guan, H., Li, F., Liu, C.Z., Dong, Y.X., Zhang, X.S. and Gao, X.Q. (2017) Arabidopsis shaker pollen inward K(+) channel SPIK functions in SnRK1 complex-regulated pollen hydration on the stigma. J Integr Plant Biol 59, 604–611.

Li, H. and Durbin, R. (2009) Fast and accurate short read alignment with Burrows-Wheeler transform. Bioinformatics 25, 1754–1760.

Li, H., Handsaker, B., Wysoker, A., Fennell, T., Ruan, J., Homer, N., Marth, G., Abecasis, G. and Durbin, R. (2009) The Sequence Alignment/Map format and SAMtools. Bioinformatics 25, 2078–2079.

Liu, D.D., Qian, W., Zhang. H.L., Fan, G.Y.Y. and Xu, Z.S. (2015) Development and Application of Molecular Markers Linked with Sex Gene X/Y in Spinach. Horticultural Plant Journal 42, 1583–1590. Chinese.

Lowe, T.M. and Eddy, S.R.J.N.a.r. (1997) tRNAscan-SE: a program for improved detection of transfer RNA genes in genomic sequence. 25, 955–964.

Luo, R.B., Liu, B.H., Xie, Y.L., Li, Z.Y., Huang, W.H., Yuan, J.Y., He, G.Z., Chen, Y.X., Pan, Q., Tang, J.B., Wu, G.X., Zhang, H., Shi, Y.J., Liu, Y., Yu, C., Wang, B., Lu, Y., Han, C.L., Cheung, D.W., Yiu, S.M., Peng, S.L., Zhu, X.Q., Liu, G.M., Liao, X.K., Li, Y.R., Yang, H.M., Wang, J., Lam, T.W. and Wang, J. (2012) SOAPdenovo2: an empirically improved memory-efficient short-read de novo assembler. GigaScience 1.

Marcais, G. and Kingsford, C. (2011) A fast, lock-free approach for efficient parallel counting of occurrences of k-mers. Bioinformatics 27, 764–770.

Murray, M.G. and Thompson, W.F. (1980) Rapid isolation of high molecular weight plant ONA. Nucleic Acids Research 8, 4321–4326.

Muyle, A., Shearn, R. and Marais, G.A. (2017) The Evolution of Sex Chromosomes and Dosage Compensation in Plants. Genome Biol Evol 9, 627–645.

Natri, H.M., Merilä, J. and Shikano, T.J.N.c. (2019) The evolution of sex determination associated with a chromosomal inversion. 10, 1–13.

Nawrocki, E.P. and Eddy, S.R.J.B. (2013) Infernal 1.1: 100-fold faster RNA homology searches. 29, 2933–2935.

Nicolas, M., Marais, G., Hykelova, V., Janousek, B., Laporte, V., Vyskot, B., Mouchiroud, D., Negrutiu, I., Charlesworth, D. and Moneger, F. (2005) A gradual process of recombination restriction in the evolutionary history of the sex chromosomes in dioecious plants. PLoS Biol 3, e4.

Okazaki, Y., Takahata, S., Hirakawa, H., Suzuki, Y. and Onodera, Y. (2019) Molecular evidence for recent divergence of X- and Y-linked gene pairs in Spinacia oleracea L. Plos One 14, e0214949.

Pannell, J.R. and Gerchen, J. (2018) Sex Determination: Sterility Genes out of Sequence. Curr Biol 28, R80–R83.

Qian, W., Fan, G., Liu, D., Zhang, H., Wang, X., Wu, J. and Xu, Z. (2017) Construction of a high-density genetic map and the X/Y sex-determining gene mapping in spinach based on large-scale markers developed by specific-locus amplified fragment sequencing (SLAF-seq). BMC Genomics 18.

Real, F.M., Haas, S.A., Franchini, P., Xiong, P., Simakov, O., Kuhl, H., Schöpflin, R., Heller, D., Moeinzadeh, M.-H. and Heinrich, V.J.S. (2020) The mole genome reveals regulatory rearrangements associated with adaptive intersexuality. 370, 208–214.

Renner, S.S. (2014) The relative and absolute frequencies of angiosperm sexual systems: dioecy, monoecy, gynodioecy, and an updated online database. Am J Bot 101, 1588–1596.

Romera-Branchat, M., Severing, E., Pocard, C., Ohr, H., Vincent, C., Née, G., Martinez-Gallegos, R., Jang, S., Andrés, F., Madrigal, P. and Coupland, G. (2020) Functional Divergence of the Arabidopsis Florigen-Interacting bZIP Transcription Factors FD and FDP. Cell Reports 31, 107717.

Stanke, M., Tzvetkova, A. and Morgenstern, B.J.G.B. (2006) AUGUSTUS at EGASP: using EST, protein and genomic alignments for improved gene prediction in the human genome. 7, S11–S11.

Starostina, E., Tamazian, G., Dobrynin, P., O’Brien, S. and Komissarov, A. (2015) Cookiecutter: a tool for kmer-based read filtering and extraction. bioRxiv.

Tennessen, J.A., Wei, N., Straub, S.C.K., Govindarajulu, R., Liston, A. and Ashman, T.L. (2018) Repeated translocation of a gene cassette drives sex-chromosome turnover in strawberries. PLoS Biol 16, e2006062.

Torres, M.F., Mathew, L.S., Ahmed, I., Al-Azwani, I.K., Krueger, R., Rivera-Nunez, D., Mohamoud, Y.A., Clark, A.G., Suhre, K. and Malek, J.A. (2018) Genus-wide sequencing supports a two-locus model for sex-determination in Phoenix. Nat Commun 9, 3969.

Turner, S.D.J.B. (2014) qqman: an R package for visualizing GWAS results using Q-Q and manhattan plots. bioRxiv.

Wadlington, W.H. and Ming, R. (2018) Development of an X-specific marker and identification of YY individuals in spinach. Theoretical and Applied Genetics 131, 1987–1994.

Wang, J., Na, J.K., Yu, Q., Gschwend, A.R., Han, J., Zeng, F., Aryal, R., VanBuren, R., Murray, J.E., Zhang, W., Navajas-Perez, R., Feltus, F.A., Lemke, C., Tong, E.J., Chen, C., Wai, C.M., Singh, R., Wang, M.L., Min, X.J., Alam, M., Charlesworth, D., Moore, P.H., Jiang, J., Paterson, A.H. and Ming, R. (2012) Sequencing papaya X and Yh chromosomes reveals molecular basis of incipient sex chromosome evolution. Proc Natl Acad Sci U S A 109, 13710–13715.

Waterhouse, R.M., Mathieu, S.,A, S.F., Mosè, M., Panagiotis, I., Guennadi, K., Kriventseva, E.V., Zdobnov, E.M.J.M.B. and Evolution (2017) BUSCO Applications from Quality Assessments to Gene Prediction and Phylogenomics. Molecular Biology & Evolution 3, 3.

Xu, C., Jiao, C., Sun, H., Cai, X., Wang, X., Ge, C., Zheng, Y., Liu, W., Sun, X., Xu, Y., Deng, J., Zhang, Z., Huang, S., Dai, S., Mou, B., Wang, Q., Fei, Z. and Wang, Q. (2017) Draft genome of spinach and transcriptome diversity of 120 Spinacia accessions. Nature Communications 8, 15275.

Yang, L., Smyth, G.K. and Wei, S.J.B. (2014) featureCounts: an efficient general purpose program for assigning sequence reads to genomic features. Bioinformatics 30, 923–930.

Yang, W., Wang, D., Li, Y., Zhang, Z., Tong, S., Li, M., Zhang, X., Zhang, L., Ren, L., Ma, X., Zhou, R., Sanderson, B.J., Keefover-Ring, K., Yin, T., Smart, L.B., Liu, J., DiFazio, S.P., Olson, M. and Ma, T. (2020) A general model to explain repeated turnovers of sex determination in the Salicaceae. Mol Biol Evol.

Yang, Z.H. (2007) PAML 4: Phylogenetic analysis by maximum likelihood. Molecular Biology & Evolution 24, 1586–1591.

Yazdi, H.P. and Ellegren, H. (2018) A Genetic Map of Ostrich Z Chromosome and the Role of Inversions in Avian Sex Chromosome Evolution. Genome Biol Evol 10, 2049–2060.

Yu, G., Wang, L.-G., Han, Y. and He, Q.-Y.J.O.a.j.o.i.b. (2012) clusterProfiler: an R package for comparing biological themes among gene clusters. OMICS: A Journal of Integrative Biology 16, 284–287.

Yu, L.a., Ma, X., Deng, B., Yue, J. and Ming, R. (2020) Construction of high-density genetic maps defined sex determination region of the Y chromosome in spinach. Molecular Genetics and Genomics.

Zhang, N., Zeng, L., Shan, H. and Ma, H. (2012) Highly conserved low-copy nuclear genes as effective markers for phylogenetic analyses in angiosperms. New Phytol 195, 923–937.

Zhao, L.N., Shen, L.K., Zhang, W.Z., Zhang, W., Wang, Y. and Wu, W.H. (2013) Ca2+-dependent protein kinase11 and 24 modulate the activity of the inward rectifying K+ channels in Arabidopsis pollen tubes. Plant Cell 25, 649–661.

