## Supplementary_info for "The female (XX) and male (YY) genomes provide insights into the sex determination mechanism in spinach": Supplementary figures.pdf

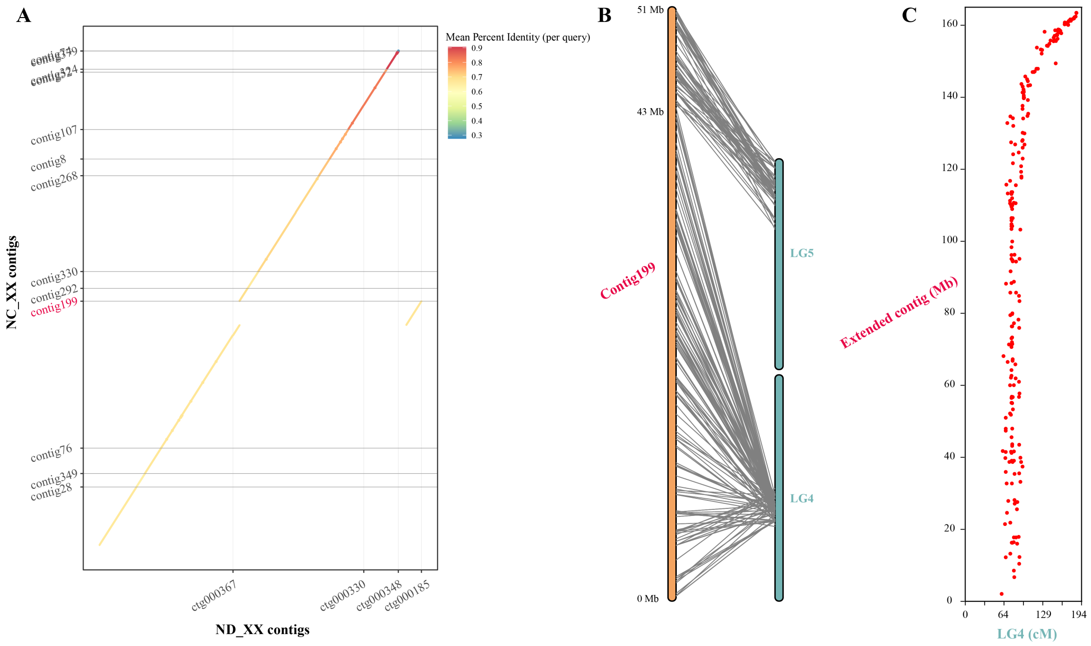


**Figure S1** An example of the correction and extension of contigs using two software programs and a linkage map. **(A)** Dot-plot alignment of some contigs from the XX genome assembly by Nextdenovo and NECAT. ND_XX contigs: the female contigs assembled by Nextdenovo; NC_XX contigs: the female contigs assembled by NECAT. **(B)** The position of SLAF markers on the contig199 and the extended contig **(C)**. LG4: linkage group 4.

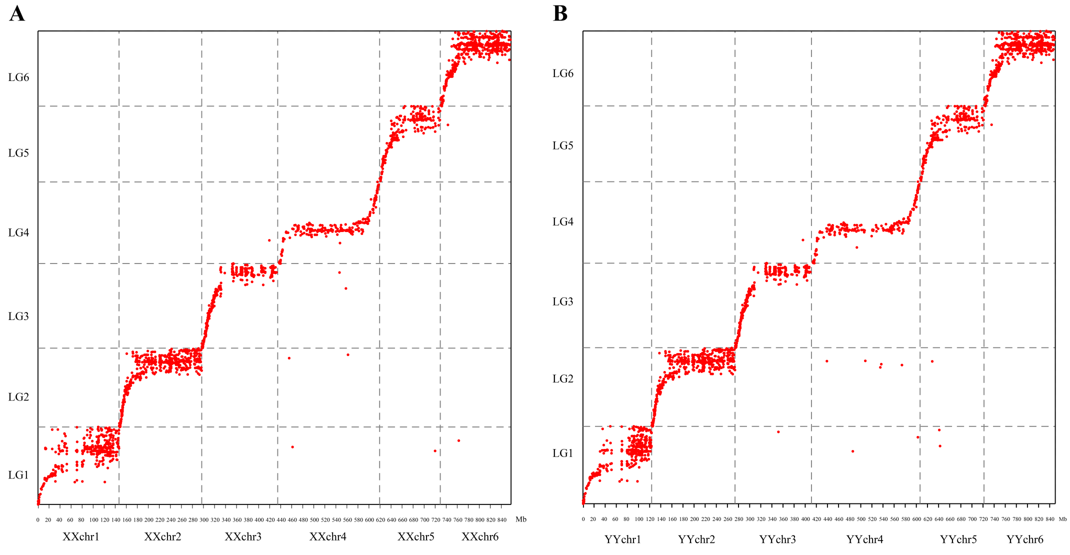


**Figure S2** Alignment of the female **(A)** and male **(B)** assembled genome with the SLAF markers linkage genetic map. LG: linkage group.

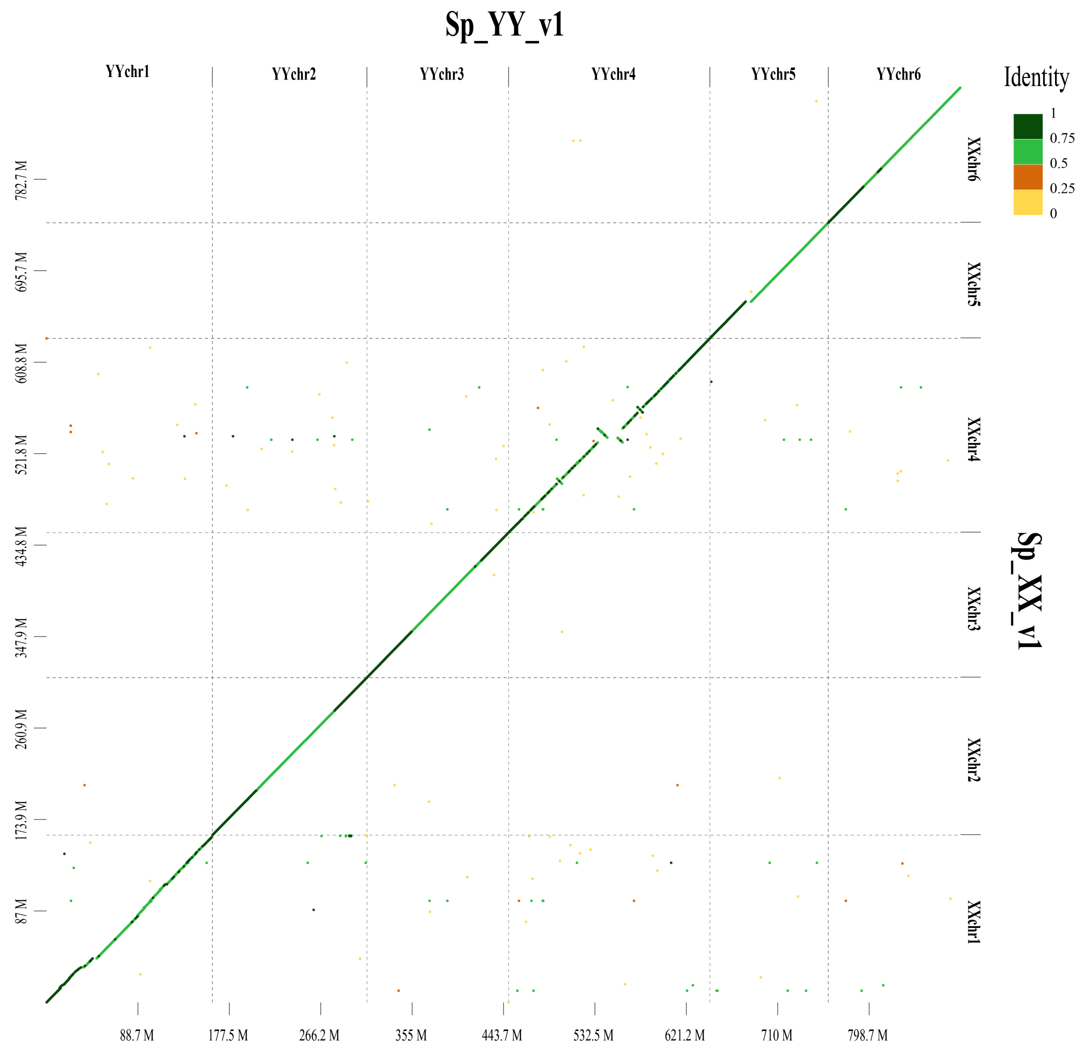


**Figure S3** Dot-plot alignment of the female (Sp_XX_v1) with male (Sp_YY_v1) genome.

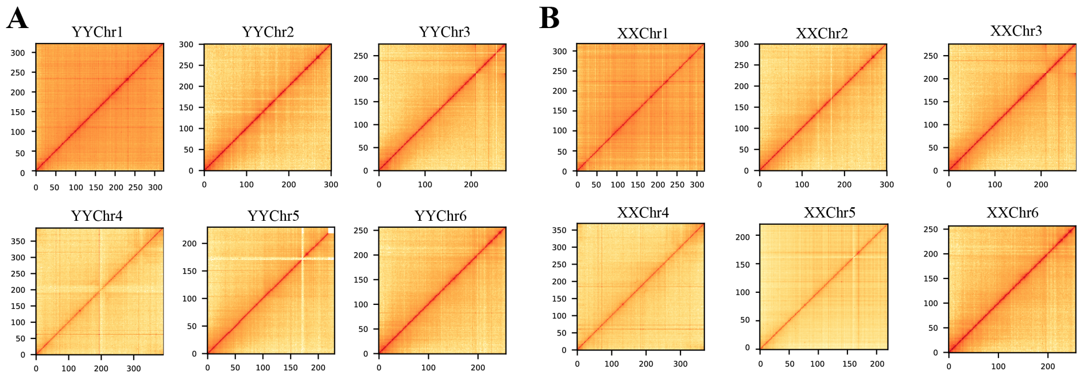


**Figure S4** Whole-genome contacts of the Hi-C data of male **(A)** and female **(B)** genomes.

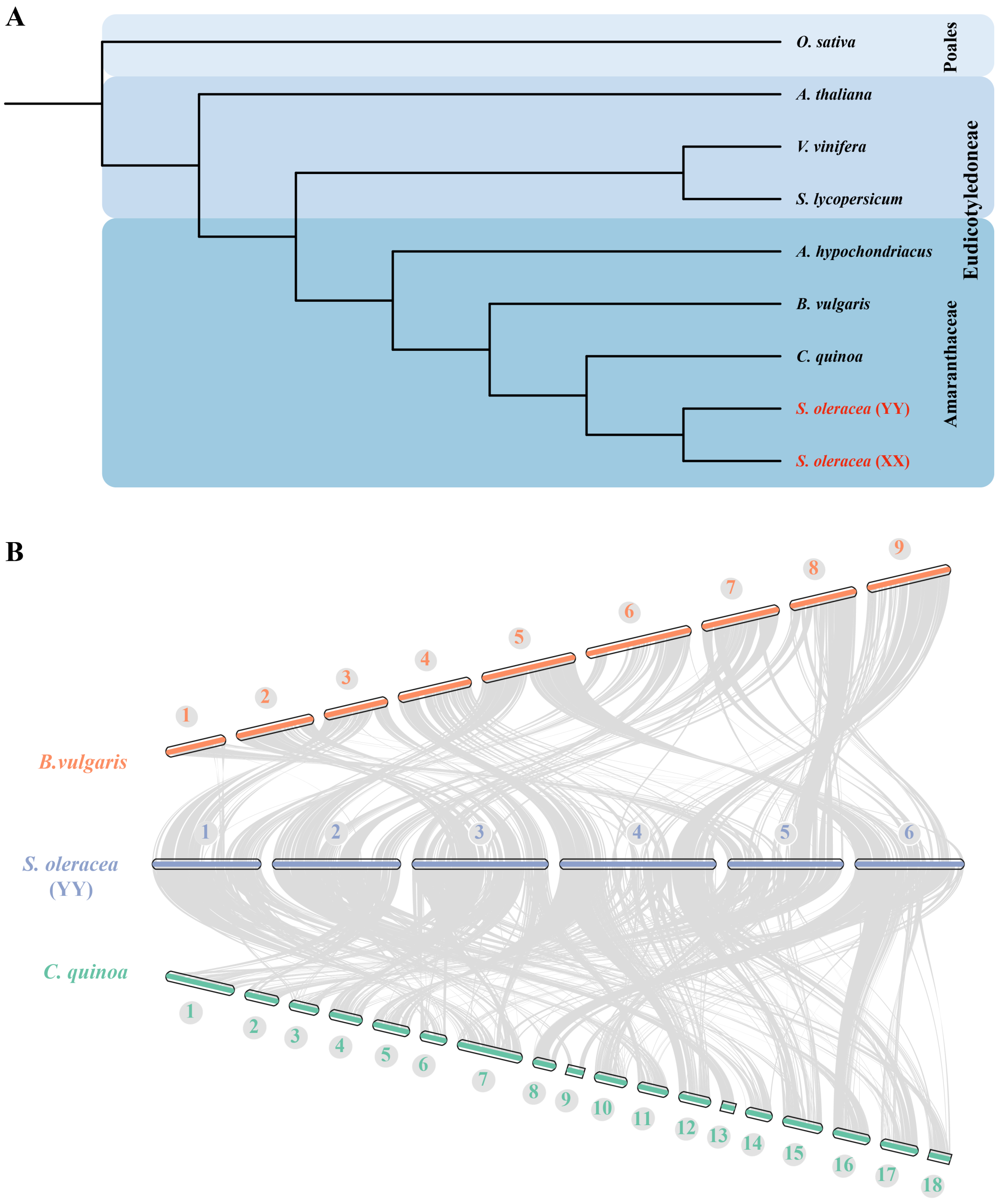


**Figure S5** Genome evolution and comparative analysis of the spinach genome. **(A)** Phylogenetic analysis of spinach and other plants. **(B)** Syntenic relationships between male (YY) spinach, sugar beet and quinoa genomes.

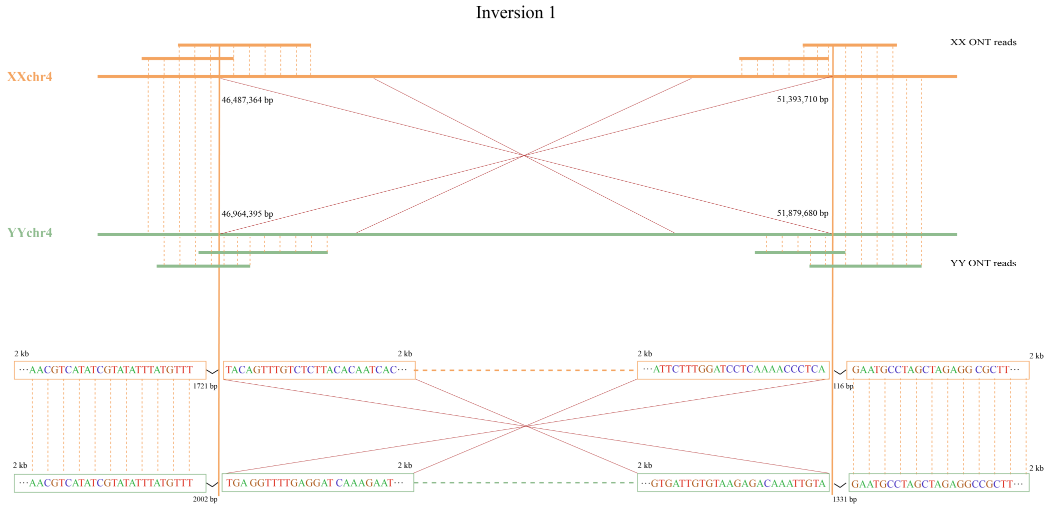


**Figure S6** Illustration of the first inversion (IV1) between the X and Y chromosome using XX and YY ONT reads, respectively. The numbers shown on the bottom (top) of the X (Y) chromosome indicate the breakpoint of the inversion.

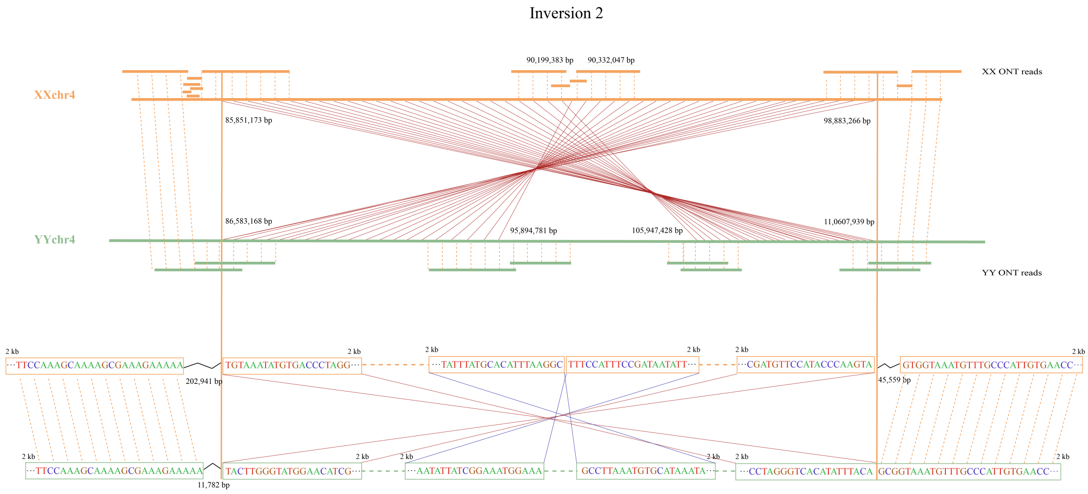


**Figure S7** Illustration of the second inversion (IV2) between the X and Y chromosome using XX and YY ONT reads, respectively. The numbers shown on the bottom (top) of the X (Y) chromosome indicate the breakpoint of the inversion.

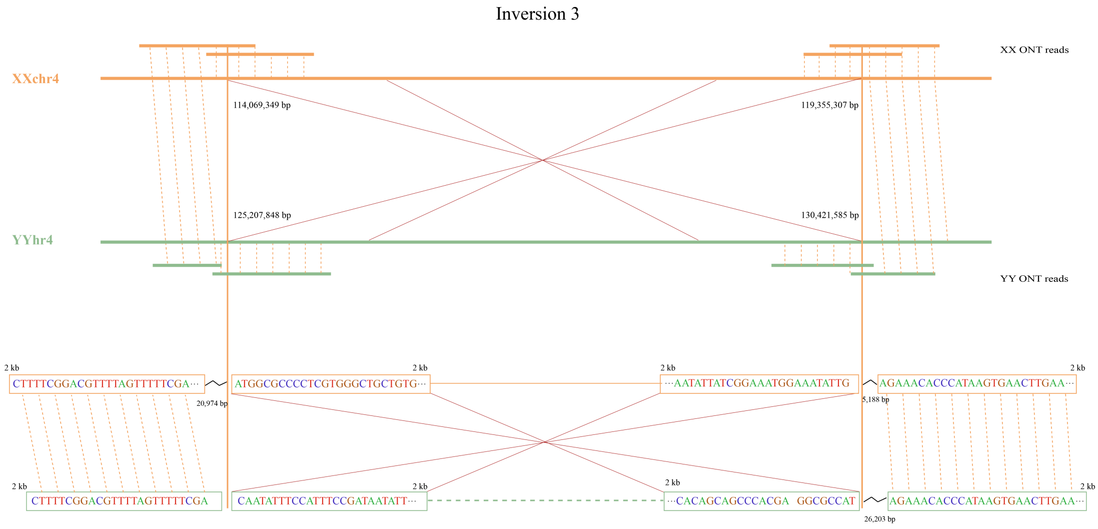


**Figure S8** Illustration of the third inversion (IV3) between the X and Y chromosome using XX and YY ONT reads, respectively. The numbers shown on the bottom (top) of the X (Y) chromosome indicate the breakpoint of the inversion.

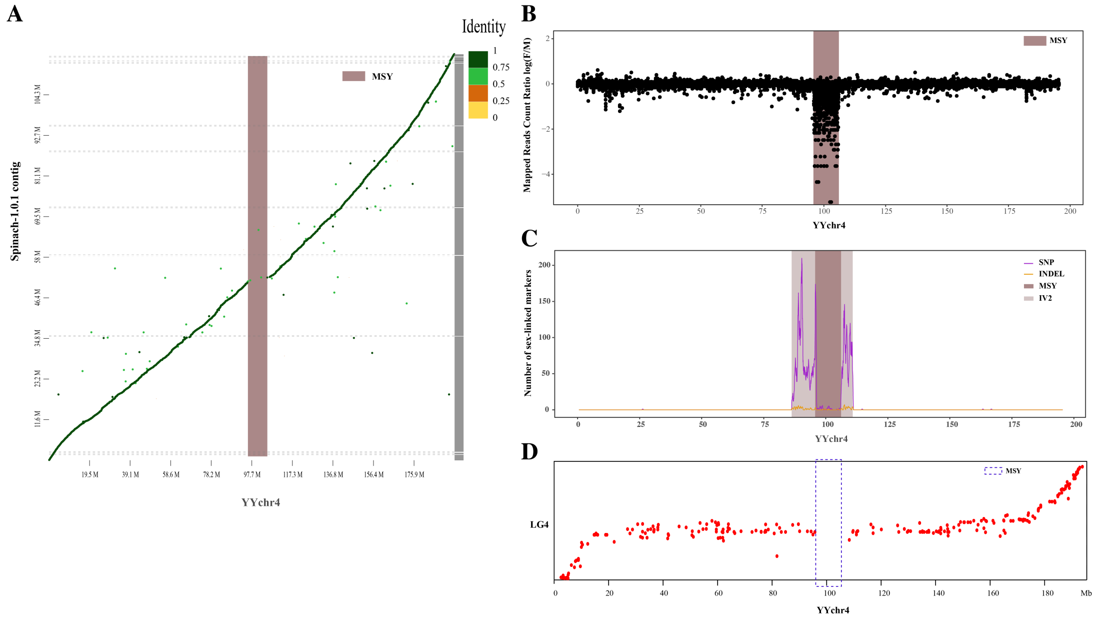


**Figure S9** Validation of the male-specific region (MSY) of Y in spinach. **(A)** Dot plot of a female genome assembled by Dohm et al. (2014) and the Y chromosome. **(B)** Sex mapped reads count ratio (log(F/M)) for 20 female and 20 male individuals are shown by black dots. The mapped reads per 10-kb bin was counted. The MSY is shaded. **(C)** Enrichment of fully sex-linked markers with 59 samples (21 female, 37 male, and one YY individuals) on Y chromosome. The number of sex-linked markers were counted for each 1 Mb. IV2 indicates the second inversion between X and Y chromosome. **(D)** The position of SLAF markers on Y chromosome. LG4: linkage group 4.

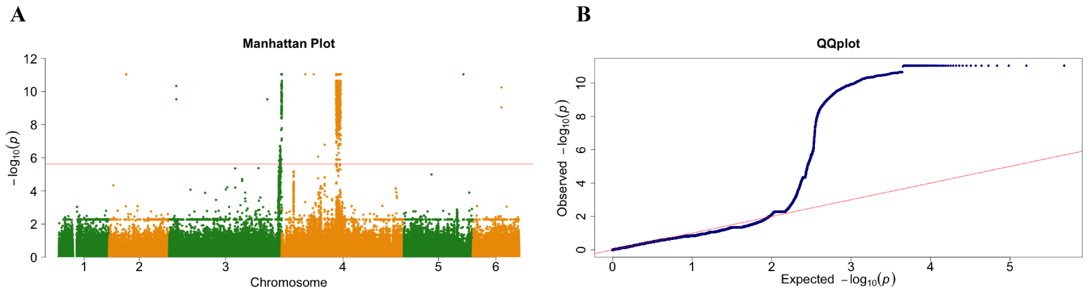


C

| ID | Position (bp) Start | End | SNPs | Female Homo (%) | Heter (%) | male Homo (%) | Heter (%) |
|---|---|---|---|---|---|---|---|
| Chr4 | 54,588,888 | 59,207,411 | 597 | 99.72 | 0.28 | 5.89 | 94.11 |
| Chr3 | 112,688,318 | 113,412,390 | 79 | 99.80 | 0.20 | 5.36 | 94.64 |
| SSC_232 | 22,201 | 2,656,358 | 250 | 99.86 | 0.14 | 5.71 | 94.29 |
| SSC_60 | 6,665 | 2,521,966 | 225 | 99.55 | 0.45 | 5.83 | 94.17 |
| SSC_66 | 2,667 | 1,722,844 | 111 | 99.27 | 0.73 | 5.12 | 94.88 |

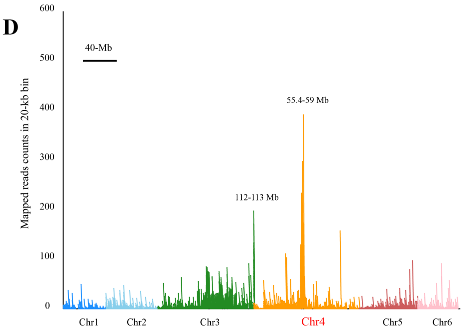


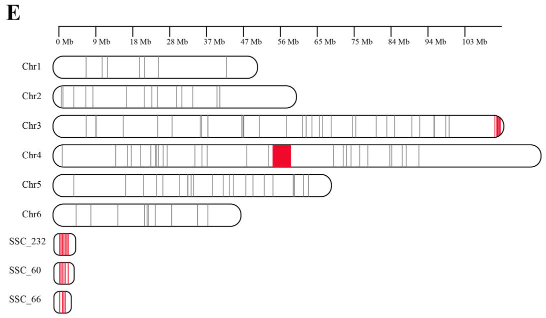


**Figure S10** Validation of the sex-determining region using GWAS and reference free k-mer. **(A)** Manhattan plot based on the results of GWAS. The y-axis represents the strength association (-$\log_{10}$($P$ value)) for each SNP. The red line indicates the threshold value ($\alpha < 0.05$). **(B)** Quantile-quantile (QQ) plot derived from genome-wide association study. **(C)** Summary of mainly SNPs significantly associated with sex. SSC: Super_scaffold, Homo: Homozygous, Heter: Heterozygosis. **(D)** Mapping of reads including male-specific k-mers (MSKs) to the spinach v1 genome. **(E)** Enrichment of sex-linked contigs on spinach v1 genome. The box indicates chromosomes/scaffolds, and the name is shown on the left. The lines within the box represent the sex-linked contigs and the sex-linked regions have been highlighted with red lines.

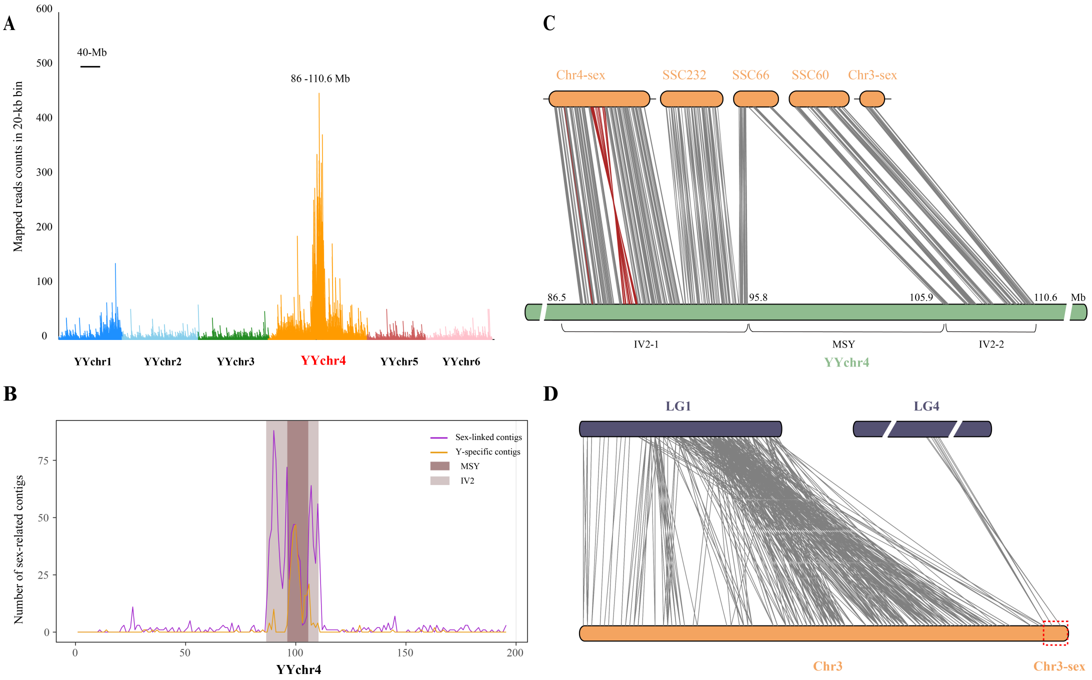


**Figure S11** Validation of the sex-related sequences identified by GWAS and k-mers approach using the male (YY) genome. **(A)** Mapping of reads including male-specific k-mers (MSKs) to the male (YY) genome. **(B)** Enrichment of sex-linked/Y-specific contigs on the Y chromosome. The 93.4% Y-specific contigs and 70.9% sex-linked contigs are enriched on IV2 and MSY. MSY: male-specific region of Y chromosome. IV2: the second inversion between the X and Y chromosome. **(C)** The position of five main sex-associated regions on the Y chromosome. Chr4-sex and Chr3-sex indicates two regions associated with sex identified in figure S10. **(D)** Alignment of the chromosome 3 from the spinach v1 genome with the SLAF markers linkage genetic map. LG: linkage group.

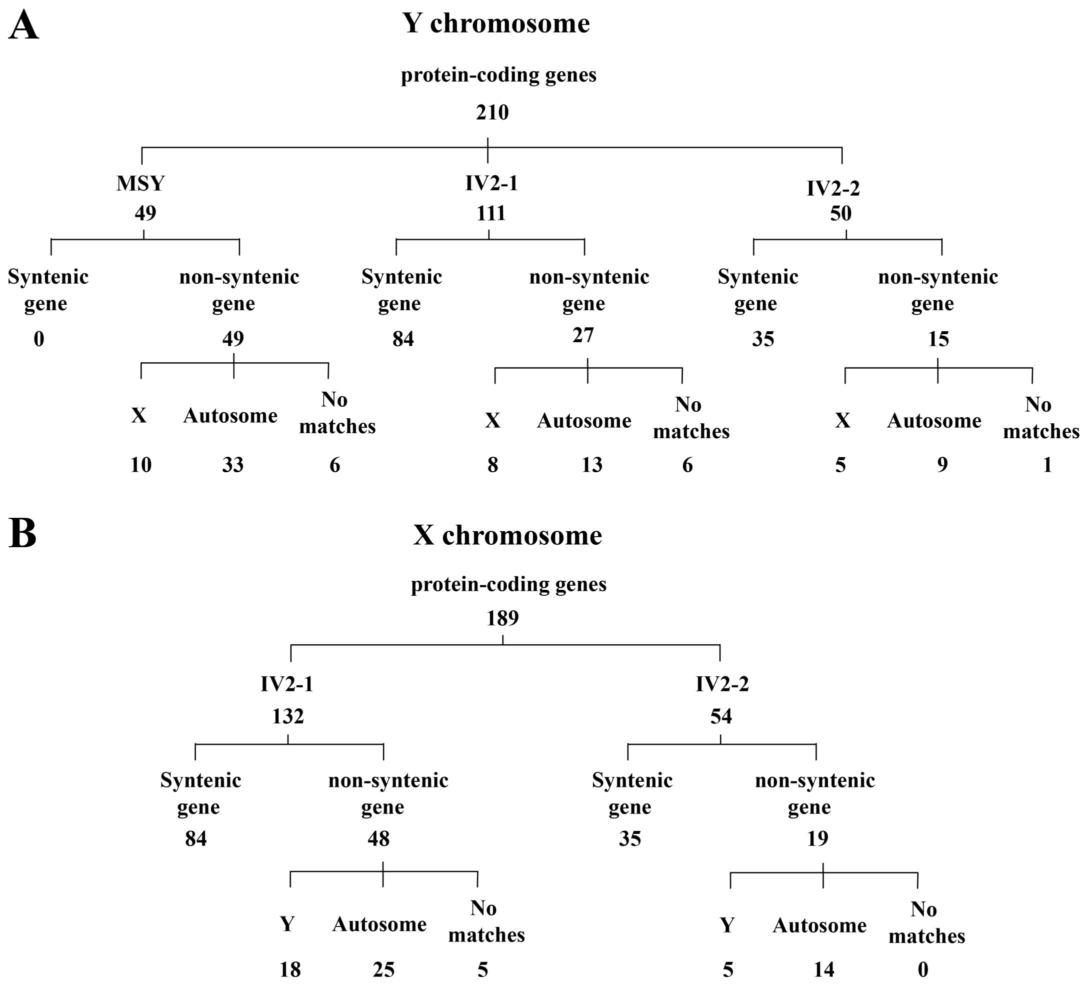


**Figure S12** Distribution of the protein-coding genes in the sex determination region of the **(A)** Y and **(B)** X chromosomes. The syntenic genes are identified using the MCscan. The orthologous genes are identified using blast with the '–evalue 1e-10'.

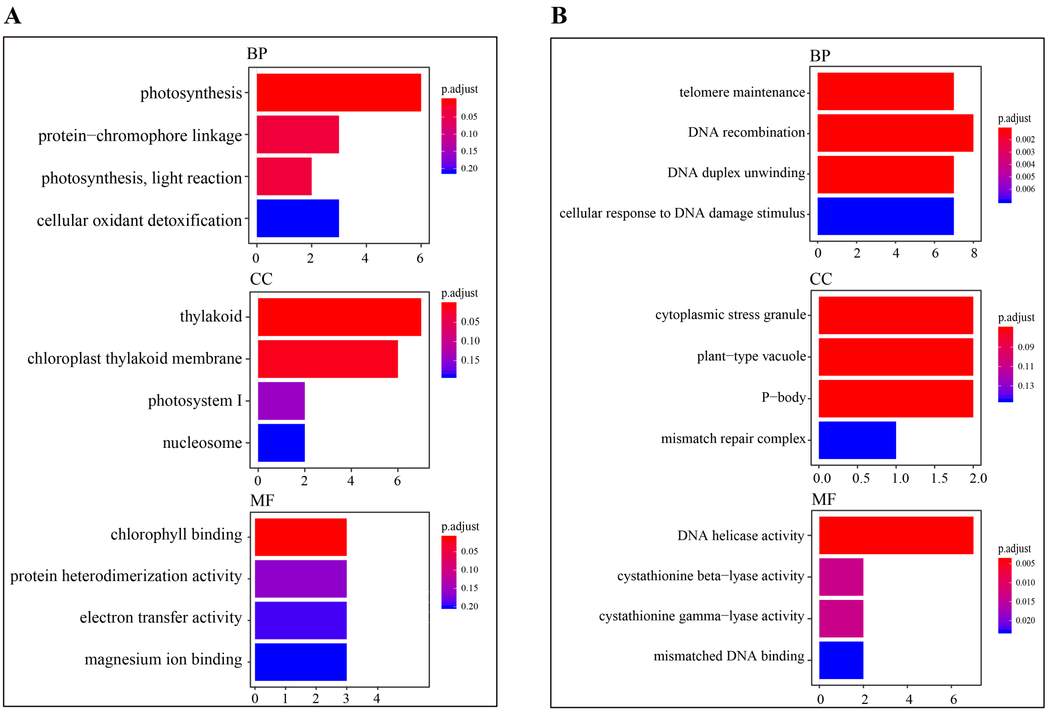


**Figure S13** GO enrichment of non-syntenic genes in sex-determining region of **(A)** the Y and **(B)** X chromosomes.

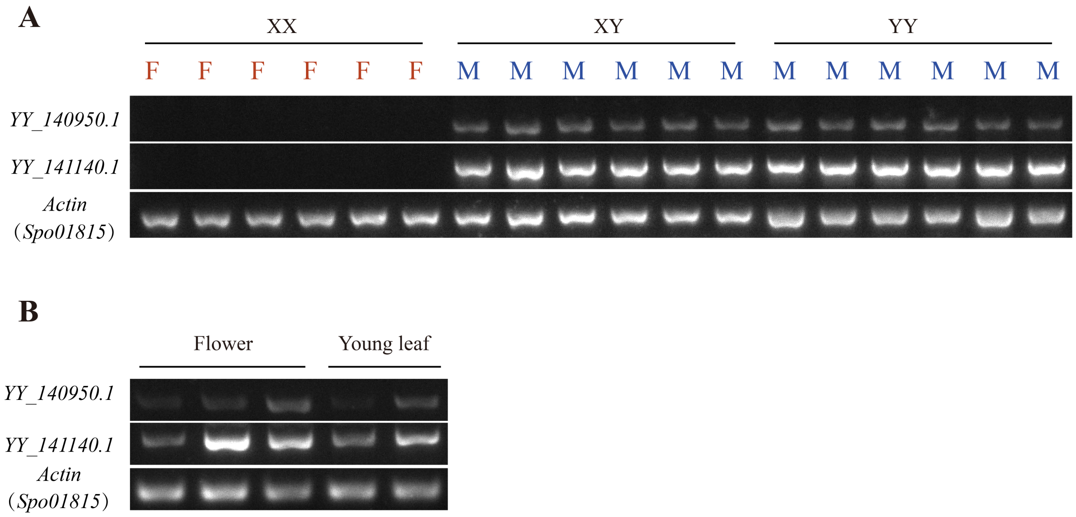


**Figure S14** Characterization of the *YY_140950* and *YY_141140* genes. **(A)** Complete male-specific conservation of *YY_140950.1* and *YY_141140.1* in the genomes of *S. oleracea*. F: female; M: male. **(B)** Expression pattern of *YY_140950.1* and *YY_141140.1* in the flower and leaf of a male individual from inbred line 10S15. The *Spo01815* is considered as an actin gene from the spinach v1 genome.

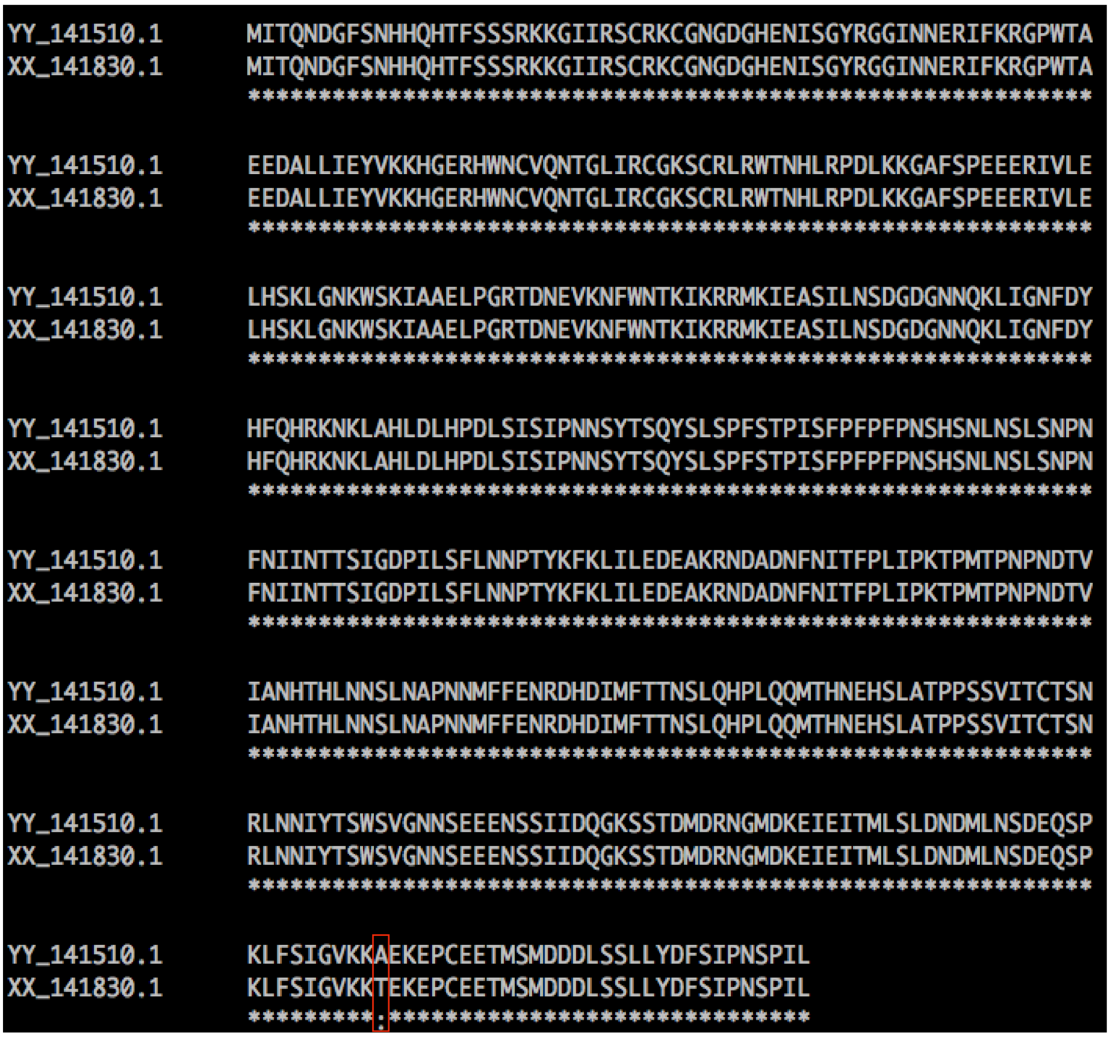


**Figure S15** Amino acid alignment of *YY_141510* and *XX_141830*. Red box indicates amino acid substitution.

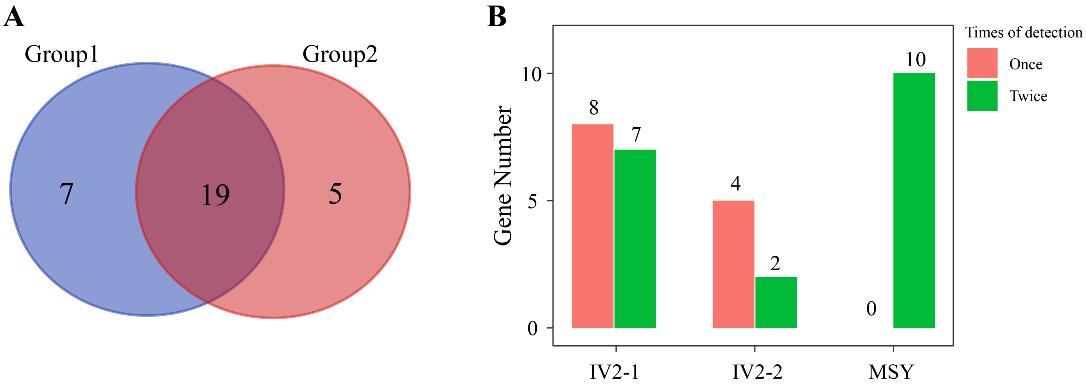


**Figure S16** Characterization of the differentially expressed genes (DEGs) identified from the sex determination region of Y chromosome. **(A)** Veen diagram of the DEGs in three groups. **(B)** Distribution of the DEGs in the IV2-1, IV2-2 and MSY.

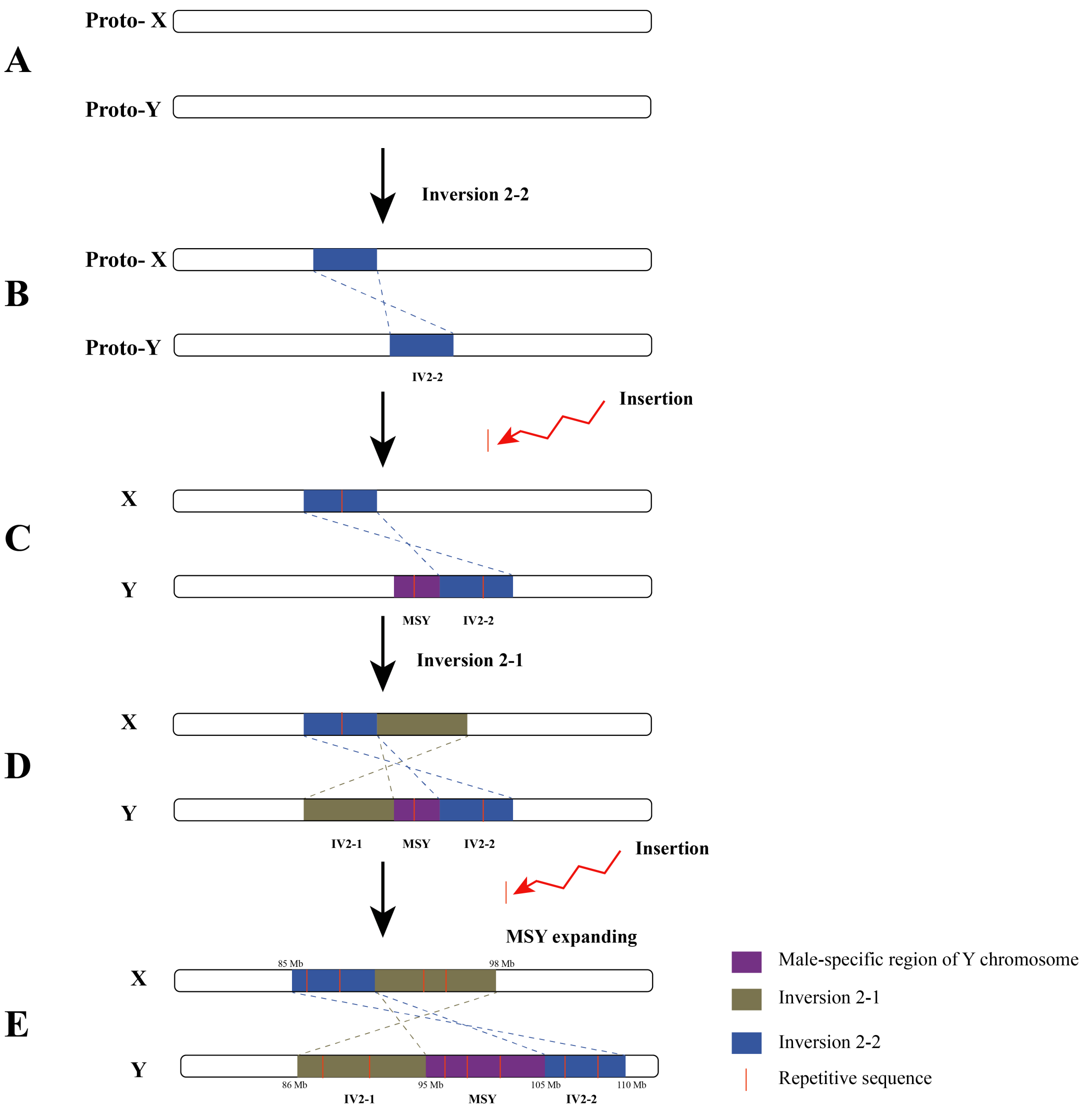


**Figure S17** Evolution of the male-specific region of Y chromosome in spinach. **(A)** The Proto-X and Proto-Y chromosome in spinach. **(B)** The IV2-2 was occurred firstly between the proto-X and proto-Y chromosome, resulting recombination suppression, repetitive sequences accumulated and generation of the MSY **(C)**. **(D)** Subsequently, the IV2-1 was occurred between X and Y chromosome, which could maintain the stableness of the MSY, thus generating larger region of the MSY **(E)**.

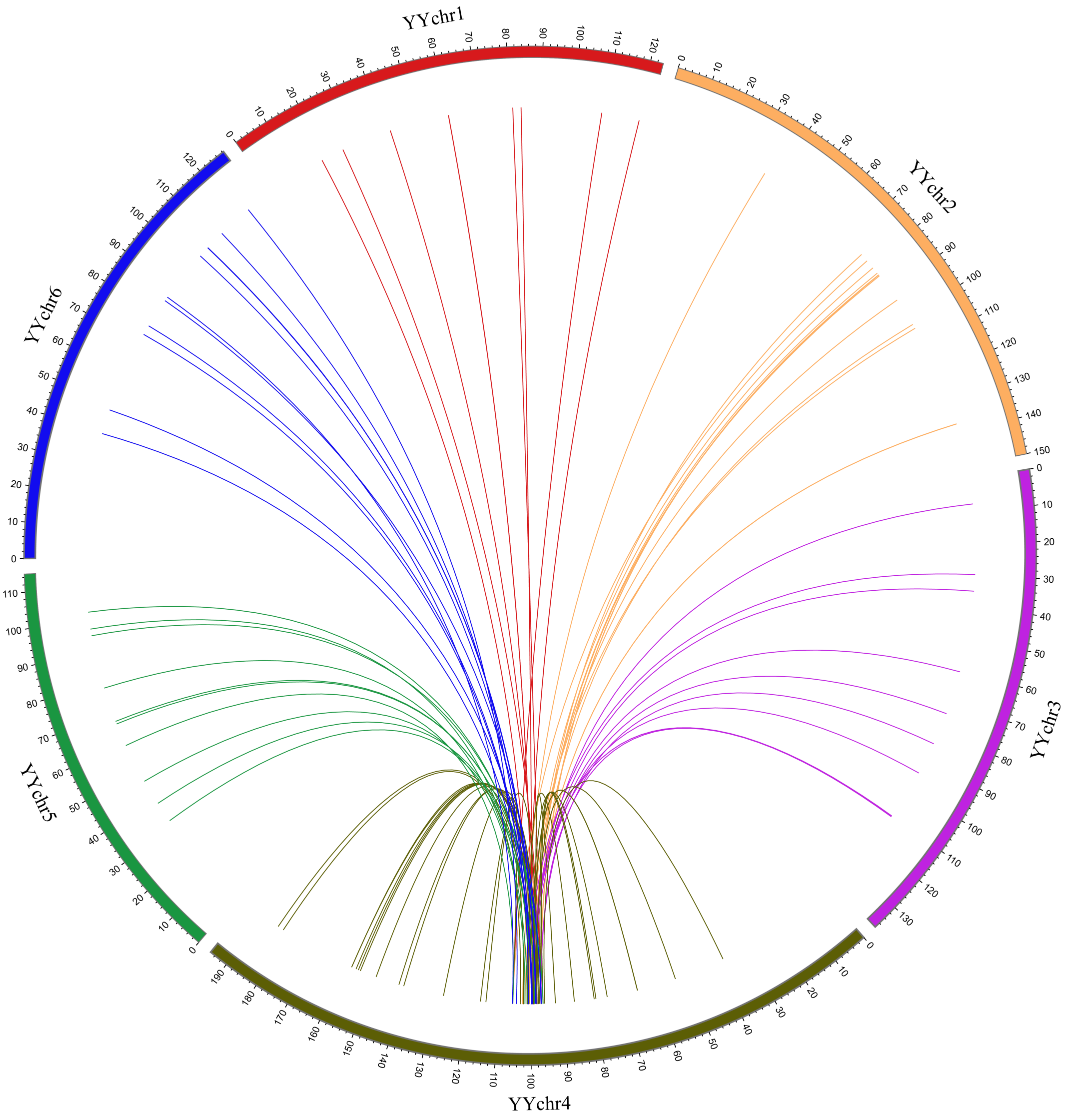


**Figure S18** Segmental duplications of the MSY and whole genome. Each line within the circus indicates similar fragment with ≥5 kb length and ≥97% identity.

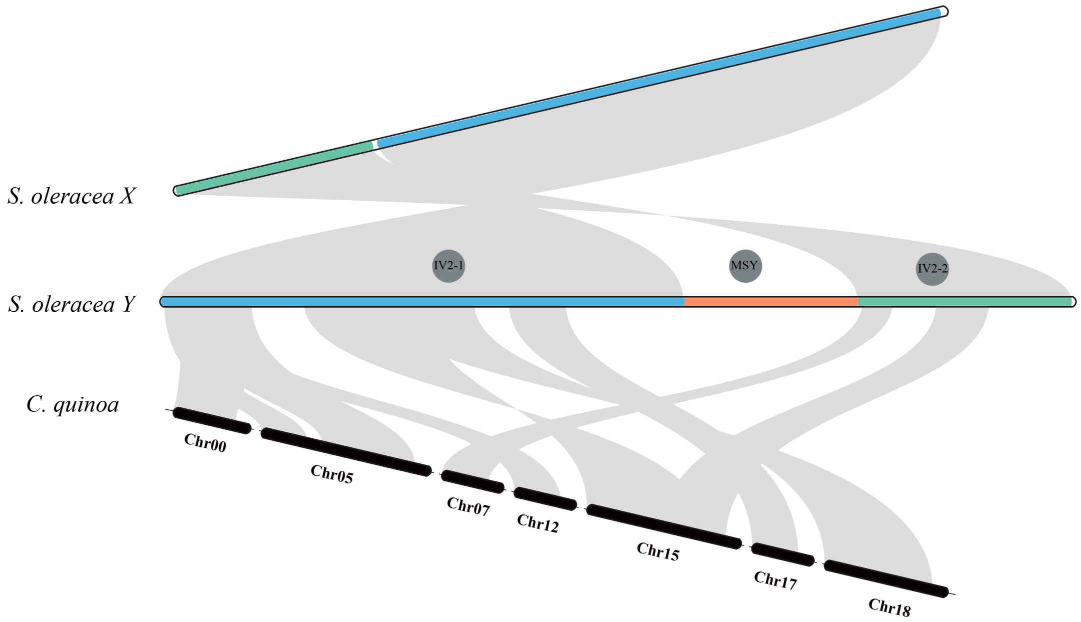


**Figure S19** Syntenic relationships between the MSY, Sp_XX_v1 and *C. quinoa* genomes. Corresponding syntenic regions are drawn. The bar with blue, orange and light green indicate IV2-1, MSY and IV2-2, respectively.
